# Gene complementation analysis suggests that dodder plants (*Cuscuta* spp.) do not depend on the host FT protein for flowering

**DOI:** 10.1101/2022.12.19.520981

**Authors:** Sina Mäckelmann, Andrea Känel, Lara M. Kösters, Peter Lyko, Dirk Prüfer, Gundula A. Noll, Susann Wicke

## Abstract

Dodder (*Cuscuta* spp.) is a genus of parasitic plants that form physiological bridges (haustoria) with their hosts to facilitate the transfer of water and nutrients. The parasites also repurpose nucleic acids and proteins translocating through the haustoria, potentially including the host florigen protein (FT), which is postulated to trigger floral transition in the parasite. Here, we identified the endogenous FT-FD flowering module in *Cuscuta campestris*. We detected the expression of two parasite-encoded *C. campestris* (Cc)FT genes in haustoria, whereas a newly found CcFD-like gene was expressed ubiquitously. *C. campestris* flowered while growing on mutant tobacco plants lacking the floral activators NtFT4 and NtFT5, indicating that host FT proteins are not required to initiate the parasite’s floral transition. We also showed that CcFT1 (identical to CaFT from *Cuscuta australis*) and CcFT2 can rescue a non-flowering *Ntft4^−^Ntft5^−^* double knockout tobacco phenotype. Together, our results show that *Cuscuta* spp. produce a potent endogenous florigen as well as other proteins likely to be involved in floral transition. FT gene expression profiles in the haustoria suggest that *Cuscuta* spp. transition to flowering at least partly in response to host signals (e.g., sugars) that can activate the parasite’s FT-FD module. Although *C. campestris* and *C. australis* appear not to depend on the host FT protein for floral transition, the nature of the mobile host signals that influence floral development in these parasites remain unclear.

**Significance Statement:** Parasitic higher plants are known for their sophisticated adaptations that facilitate the transfer of water and nutrients from their hosts. They can also synchronize their transition from vegetative to reproductive development to match the host plant. Despite this high degree of synchronization, dodder plants maintain a potent endogenous floral activator module, which enables the parasite to switch to reproductive development autonomously. Synchronization must therefore involve other stimuli from the host plant, which are currently unknown. Understanding the environmental cues that trigger flowering, and the corresponding network of genetic and physiological regulators and integrators, may lead to new strategies that reduce the reproductive fitness of parasitic plants to protect crops and ensure food security.

**Data Servers:** This article is available as preprint (ID: BIORXIV/2022/520981) at https://www.biorxiv.org under the CC BY-NC 4.0 license. Reusable data files have been deposited at https://datadryad.org, accessible during peer-review under: https://datadryad.org/stash/share/DK8Olh2VqFwbGNL0GtkGt24dD0GhWhJn82oLBC1XK70

## Introduction

*Cuscuta* is a genus of ~4 500 parasitic plants that survive by taking water and nutrients from their hosts (1). The parasites can survive a few days after germination but must find a host plant to complete their life cycle. *Cuscuta* spp. have a broad host range, including many cultivated crops, and are therefore a threat to global agriculture. *Cuscuta* plants form specialized feeding organs (haustoria) that connect to the host vascular tissue, through which they acquire water, nutrients, secondary metabolites, small RNAs, mRNAs, and proteins (2–5). Among the latter is the florigen FT, a peptide hormone, which was found to translocate from host to parasite and was recently proposed to cause the synchronization of flowering in the parasite–host system (6).

Flowering is tightly controlled by endogenous and environmental cues, including hormones, plant age, temperature, and day length, to ensure that floral development occurs under favorable conditions. It is best studied in the winter-annual *Arabidopsis thaliana*, which is regarded as the principal model of flower development. In this species, the transcription factor FLOWERING LOCUS C (FLC) represses flowering by downregulating the floral integrator genes *FLOWERING LOCUS T (FT), SUPPRESSOR OF OVEREXPRESSION OF CONSTANS 1 (SOC*), and *FD*, the latter encoding the cofactor of FT (7–10). A prolonged period of cold (vernalization) reduces *FLC* mRNA levels by inducing the persistent methylation of histones at the *FLC* locus (11, 12). The corresponding abolition of FLC activity is a key milestone in floral induction. Day length also regulates flowering in Arabidopsis because the CONSTANS (CO) protein is only stable under long-day (LD) conditions, and a stable CO protein is required to activate *FT* gene expression (13). FT is expressed mainly in the leaves, but it is transported through phloem sieve elements to the shoot apical meristem (SAM), where it interacts with its cofactor FD and 14-3-3 proteins, together forming the floral activating complex (FAC) (14, 15). This protein complex enhances the expression of *SOC1* and induces the expression of the floral meristem identity gene *APETALA1 (AP1*) (16). Meanwhile, SOC1 interacts with the MADS transcription factor AGAMOUS-like 24 (AGL24) to upregulate the expression of *LEAFY (LFY*) (17). LFY and AP1 then promote the formation of floral meristems, the first step in the process of flower development (18, 19).

Many plant species differ from Arabidopsis and follow a diverse set of floral initiation strategies (20). The parasite *Cuscuta australis* was proposed to require host FT proteins to complement its own, non-functional FT gene to allow floral induction (6). Genome sequencing of two *Cuscuta* species revealed varying degrees of gene loss compared with autotrophic plants (21, 22). For *C. australis*, the loss of 26 of the 295 proteincoding genes in the Arabidopsis flowering network database was postulated based on bioinformatic analysis (22, 23). The lost genes included *CO* and *FLC*, suggesting *C. australis* has a unique strategy to regulate flowering time (22). The eventual discovery of host FT proteins in *C. australis* tissue led researchers to conclude that the host FT and endogenous FD form a complex to enable floral transition in the parasite (6). However, this contrasts with the observation that *Cuscuta* spp. flower independently *in vitro* (24, 25). Furthermore, we have never observed any impact of the host’s flowering status on the flowering time in *Cuscuta campestris*, the model parasitic plant studied in our laboratory. We therefore investigated the ability of *C. campestris* to flower independently by allowing it to parasitize wildtype tobacco and CRISPR/Cas9 mutants lacking FT floral activators. We also examined the expression of *CcFT* mRNAs in different tissues and their ability to complement tobacco FT mutants. Our results provide insight into the regulation of flowering in *Cuscuta* spp., and suggest that it is not dependent on host FT proteins.

## Results

### Host FT expression does not affect flowering of *C. campestris*

We analyzed the flowering of *C. campestris* plants parasitizing wild-type SR1 tobacco and a non-flowering mutant. Tobacco is a day-neutral plant that produces two FT floral inducers (NtFT4 and NtFT5). NtFT5 is predominantly responsible for floral induction under LD conditions, whereas both NtFT4 and NtFT5 induce flowering under short-day (SD) conditions (26, 27).

We transformed a well-characterized and stable *Ntft5^−^* mutant (28) with a CRISPR/Cas9 construct to knock out *NtFT4*. Following the regeneration of double knockout plants homozygous for mutations in *NtFT4* and *NtFT5* (*Ntft4^−^Ntft5^−^*; Fig. S1), the shoots of the mutant plants were grafted onto wild-type tobacco stocks to enable flowering and seed formation. The *Ntft4^−^Ntft5^−^* genotype was confirmed in the offspring, and phenotypic analysis revealed that T_1_ mutant plants were unable to flower under LD or SD conditions up to the end of the experiment (139 and 120 days, respectively). In contrast, wild-type SR1 tobacco flowered after 62–64 days under LD or SD conditions (Fig. S2).

We allowed *C. campestris* to parasitize wild-type tobacco plants and two independent *Ntft4^−^Ntft5^−^* lines, and we monitored the flowering behavior under LD conditions. The wild-type SR1 plants flowered after 66 days, whereas the *Ntft4^−^Ntft5^−^* plants had not flowered by the end of the experiment (90 days). *C. campestris* plants parasitizing wild-type tobacco flowered after 42 days, and those parasitizing the mutant tobacco plants flowered after 39 and 42 days, respectively (Fig. 1). These data suggest that *C. campestris* does not rely on the host FT signal for the transition to flowering.

**Fig. 1.**
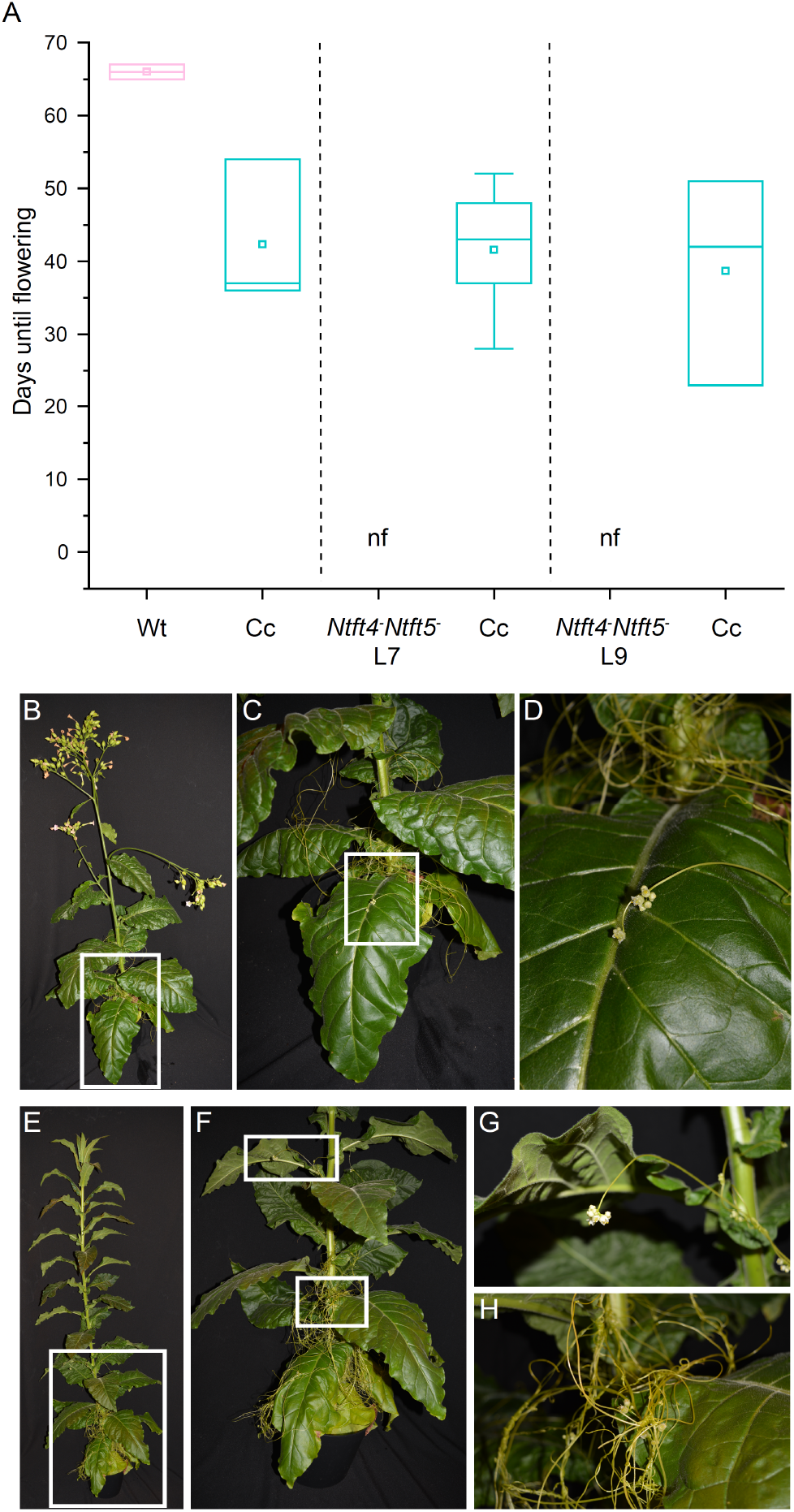
*C. campestris* parasitizing tobacco with varying flowering habits. **A)** *C. campestris* (Cc) was allowed to parasitize tobacco SR1 wild-type plants (Wt) or nullizygous *Ntft4^−^Ntft5^−^* plants representing two independent transformation events (lines 7 and 9). Days from seeding (tobacco) or haustorium formation (*C. campestris*) until flowering were determined when the first bud fully opened. No significant differences in flowering time were observed between *C. campestris* plants parasitizing flowering and nonflowering tobacco hosts as determined by ANOVA and Tukey’s *post hoc* test (*n* ≥ 3 biological replicates); nf – non-flowering until the end of experiment (83 days after sowing). In the boxplots, center line = median, square = mean, box limits = upper and lower quartiles, and whiskers = 1.5× interquartile range. **B-H)** Flowering phenotypes of *C. campestris* parasitizing SR1 wild-type **(B-D)** or non-flowering *Ntft4^−^Ntft5^−^* **(E-H)** plants. Magnified areas from B (C, D) and E (F, G, H) are indicated by white squares. Photographs were taken 83 days after sowing.

### Identification of endogenous *FT* and *FD* genes in *Cuscuta* spp

The endogenous FT-FD module in *C. campestris* was examined in detail using genomic and transcriptomic data of *Cuscuta* spp. in combination with gene family analyses (Tables S1, S2, S3). Multispecies analysis of 295 flowering time-associated protein-coding genes indicated only two putative *Cuscuta*-specific gene deletions (UBIQUITIN-SPECIFIC PROTEASE 26; GIBBERELLIN 2-OXIDASE 8; Figure S3), whereas other gene losses (e.g., ELF4) or considerable divergence from validated coding sequences of *Arabidopsis* are shared with other eudicots (Fig. S3; Supplemental Dataset 1). We amplified and sequenced the complete genomic sequences of *CcFT1* (11,502 bp) and *CcFT2* (12,819 bp). The corresponding coding sequences were 534 and 531 bp in length, respectively. Despite pseudogenized domains of a non-intact *Ty1/copia Ale-type* retrotransposon in the intron separating exons 3 and 4 of *CcFT2* and a LINE-type reverse transcriptase upstream of *CcFT1* and the *C. australis* orthologue CaFT (Figure S4; Supplemental Dataset 2), all *FT* genes splice correctly according to RT-PCRs results. Phylogenetic analysis showed that CcFT1, CcFT2, and CaFT cluster with FT homologs of the phosphatidylethanolamine-binding protein (PEBP) family from other species (Fig. S5). The deduced amino acid sequences of CcFT1 and CaFT were identical (Fig. S6). To identify the *C. campestris* homolog of FD, we screened genomic and transcriptomic data using *FD* genes of *Nicotiana tabacum, Arabidopsis thaliana, and Ipomoea nil* as queries. We identified seven and five new *FD*-like sequences in *C. australis* and *C. campestris*, respectively, whereby we paid particular attention to a C-terminal SAP motif that distinguishes FD proteins from other bZIP transcription factors. The association of FDs with 14-3-3 proteins and their interaction with FTs depends on the phosphorylation of threonine or serine residues in this SAP motif (8, 29). Only two of the identified transcripts of *C. campestris* encoded an SAP-like motif, herein named *CcFD-like1* and *CcFD-like2* (Fig. S7 and S8). The corresponding amino acid sequences were 99% identical. We also identified a sequence identical to *CcFD-like1* in the genome of *C. australis* (*CaFD-like*). This new *CaFD-like* gene differs from the previously published *CaFD* sequence, whose protein product lacks the C-terminal SAP motif and was reported to not interact with *C. australis*’ endogenous FT protein (6).

### *C. campestris* and *C. australis* express endogenous *FT* and *FD* mRNAs

We analyzed the expression of *CcFT* and *CcFD-like* genes by quantitative real-time PCR (qRT-PCR) using total RNA isolated from stems, stem tips, branches, scale-like leaves, haustoria, buds, and flowers of flowering *C. campestris* plants parasitizing wild-type tobacco (Fig. S9; Table S4). We detected *CcFD-like* expression in all tissues, similarly strong in haustoria, buds and flowers, but slightly weaker in the other tissues. We detected *CcFT1* and *CcFT2* expression only in haustoria, with low expression levels in both cases (Fig. 2A). The integrity of the amplicons was verified by Sanger sequencing. Because residual tobacco tissues may have been co-harvested with the *C. campestris* haustoria, we also tested tobacco cDNA with the same primer sets (Table S4). We did not detect any amplicons, confirming the primer specificity (Fig. 2A). Next, we repeated the qRT-PCR assay using total RNA from *C. australis* plants parasitizing soybean (*Glycine max*). Similarly, we detected *CaFD-like* gene expression in all tissues, whereas *CaFT* was haustorium-specific (Fig. 2B). We then identified the 3’-UTRs of the *FT* mRNAs by 3’-RACE, allowing us to amplify the complete coding sequences, including parts of the 3’-UTR of *CcFT1, CcFT2* and *CaFT*. These results were further confirmed by RNA-Seq data analysis of FT and FD genes, alongside other core flowering time regulators, which we detected in both *Cuscuta* species (Tables 1 and S1). Together, these results show that *C. campestris* express two paralogs each of FT and its cofactor FD, as well as flowering time regulators interacting with these. In contrast to previous findings (6), we also detected *CaFT* and *CaFD*-like gene expression in *C. australis*.

**Fig. 2.**
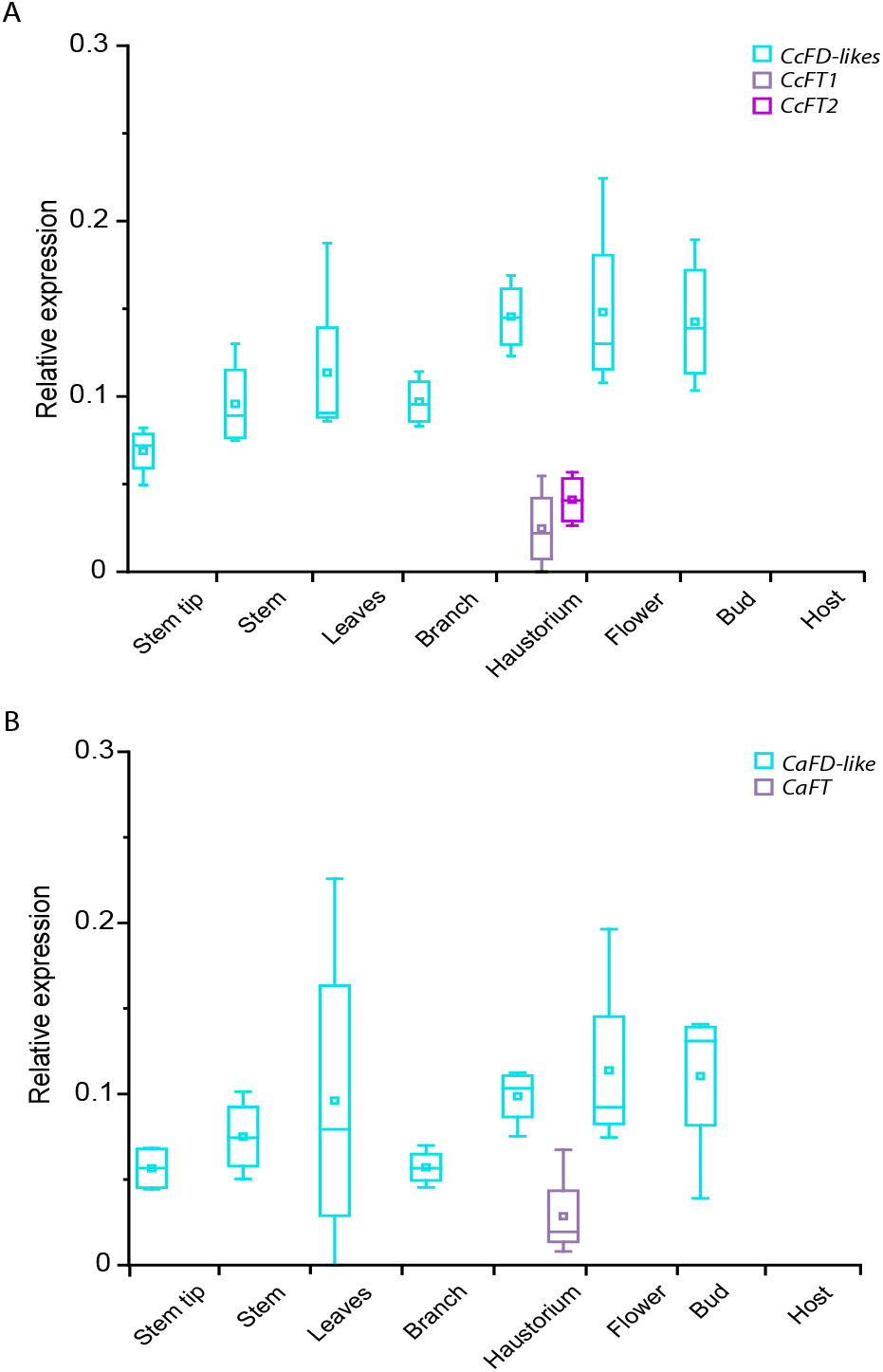
Expression of *FD* and *FT* homologs in *Cuscuta* spp. **A)** Relative expression levels of *CcFT1, CcFT2* and *CcFD*-like genes were determined by qRT-PCR, using *Actin* and *EF1α* as reference genes (*n* = 4 biological replicates). Plant material was harvested from flowering *C. campestris* plants parasitizing wildtype tobacco SR1. **B)** Relative expression levels of *CaFT* and *CaFD*-like were determined by qRT-PCR using *Actin* and *EF1α* as reference genes (*n* = 4 biological replicates). Plant material was harvested from flowering *C. australis* plants parasitizing soybean. In the boxplots, center line = median, square = mean, box limits = upper and lower quartiles, and whiskers = 1.5× interquartile range.

**Table 1.** RNA Seq-based transcript levels.

|  | Gene length | *Cuscuta campestris*<br>total reads mapped:<br>1 990 329 102 (after trim) |  | *Cuscuta australis*<br>total reads mapped:<br>908 853 342 (after trim) |  |
| --- | --- | --- | --- | --- | --- |
|  |  | Coverage | RPKM | Coverage | RPKM |
| FT1 | 534 bp | 4.56 | 57.01 | 12.98 | 633.80 |
| FT2<sup>1</sup> | 531 bp | 4.68 | 64.81 | -- | -- |
| FD-like | 918 bp | 1 025.49 | 11 596.03 | 398.02 | 16 046.96 |
| SOC1 | 714 bp | 43 406.26 | 498 893.10 | 6 731.04 | 297 048.13 |
| TFL1 | 594 bp | 1 724.57 | 22 286.12 | 5 020.35 | 226 723.48 |
| LFY | 1263 bp | 236.22 | 2 592.57 | 313.55 | 13 852.78 |
| CO-like | 1047 bp | 3 986.97 | 50 309.32 | 2 676.82 | 123 982.11 |
| GI-like | 3453 bp | 8 752.11 | 97 400.03 | 1 981.41 | 79 456.96 |

<sup>1</sup> not found in *C. australis*

### Interactions between *Cuscuta* spp. FT and FD-like proteins

FT interacts with FD to form the FAC. We studied the interaction between CcFT1/CcFT2 and CcFD-like1 by bimolecular fluorescence complementation (BIFC) based on the monomeric red fluorescent protein (mRFP1). Confocal laser scanning microscopy revealed that both CcFT1 and CcFT2 interact with CcFD-like1 in the nucleus of *N. benthamiana* epidermal cells in transient expression experiments (Fig. S10). CcFT1 and CcFT2 also interacted with NtFD1, and CcFD-like1 was able to interact with NtFT5 (Fig. S10). These results highlight the conservation of the FT-FD module across species, as shown for other interspecific pairs (26, 30). The amino acid sequences of CcFT1 and CaFT, as well as CcFD-like1 and CaFD-like, are identical, and all the corresponding mRNAs are expressed. In sum, these results indicate that an endogenous FT-FD protein complex can form in both *C. campestris* and *C. australis*.

### Heterologous expression of *Cuscuta* FTs enables the flowering of non-flowering mutants

Despite significant progress toward the genetic transformation of *Cuscuta reflexa* (31), a protocol for the generation of stably transformed *Cuscuta* plants is not yet available. This genus is phylogenetically closely related to tobacco, which is more amenable to genetic modification, so we expressed the *CcFT1* gene in tobacco using the strong and constitutive CaMV 35S promoter. We analyzed the T_1_ progeny of three independent transgenic lines, revealing that CcFT1 overexpression did not strongly affect flowering time. One of the three lines showed a slight but significant increase in flowering time (~4.5 days) compared with vector controls (Fig. S11). Given that CcFT1 overexpression fails to induce early flowering in tobacco, it appears that CcFT1 cannot surpass the function of the two endogenous activators, NtFT4 and NtFT5. Next, we tested whether CcFT1 or CcFT2 was able to complement the non-flowering phenotype of the *Ntft4^−^Ntft5^−^* double knockout plants. The regenerated T_0_ plants were indeed able to flower spontaneously, allowing seeds to be harvested. For phenotypic analysis of the T_1_ progeny, we chose three lines per construct with strong expression levels in the T_0_ generation. Under LD conditions, *Ntft4^−^Ntft5^−^* plants overexpressing *CcFT1* flowered 67–207 days after seed sowing (Fig. 3). Only two of 48 plants had not flowered after 233 days (end of experiment), probably due to their low *CcFT1* expression levels (Fig. S12). *Ntft4^−^Ntft5^−^* plants overexpressing *CcFT2* flowered 70–130 days after sowing, whereas the *Ntft4^−^Ntft5^−^* internal control plants had not flowered after 233 days (Fig. 3). These results show that CcFT1 (and by extension the identical CaFT) and CcFT2 can induce floral transition in a heterologous system and should therefore be considered functional floral activators. This, together with their endogenous expression and confirmed interaction with CcFD-like1 and CaFD-like, respectively, strongly indicates that floral induction at least in these two *Cuscuta* species is not dependent on FT proteins imported from host plants.

**Fig. 3.**
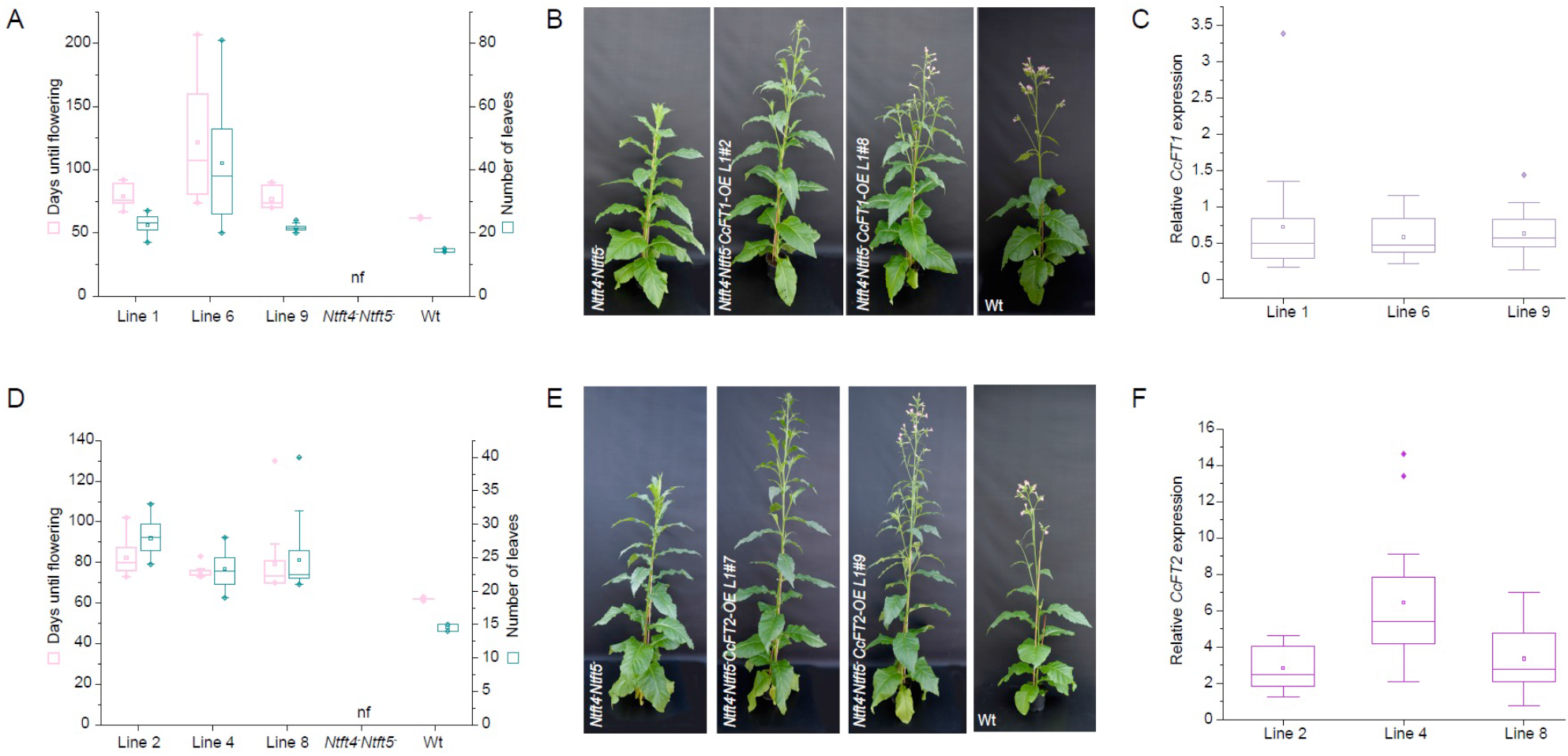
Overexpression of *CcFT1* or *CcFT2* in *Ntft4^−^Ntft5^−^* tobacco plants. **A)** Flowering time and number of leaves at flowering of *Ntft4^−^Ntft5^−^* plants overexpressing *CcFT1* compared to *Ntft4^−^Ntft5^−^* and wild-type (Wt) controls. Days until flowering and number of leaves were determined when the first bud had fully opened; *n* = 17 (line 1), *n* = 15 (line 6), *n* = 14 (line 9), *n* = 12 (*Ntft4^−^Ntft5^−^*), *n* = 6 (Wt). **B)** Phenotypes of *Ntft4^−^Ntft5^−^* plants with or without the heterologous expression of *CcFT1* compared to Wt controls at the time of Wt plants flowering. **C)** Relative expression levels of *CcFT1* in *Ntft4^−^Ntft5^−^* plants overexpressing *CcFT1*. **D)** Flowering time and number of leaves at flowering of *Ntft4^−^Ntft5^−^* plants overexpressing *CcF2* compared to *Ntft4^−^Ntft5^−^* and Wt controls. Days until flowering and number of leaves were determined when the first bud had fully opened; *n* = 12 (line 2), *n* = 17 (line 8), *n* = 18 (line 9), *n* = 13 (*Ntft4^−^Ntft5^−^*), *n* = 6 (Wt). **E)** Phenotypes of *Ntft4^−^Ntft5^−^* plants with or without the heterologous expression of *CcFT2* compared to Wt controls. Pictures were taken when Wt plants flowered. **F)** Relative expression levels of *CcFT2* in *Ntft4^−^Ntft5^−^* plants overexpressing *CcFT2*; nf – non-flowering until end of experiment (233 days after sowing). In the boxplots, center line = median, square = mean, box limits = upper and lower quartiles, whiskers = 1.5× interquartile range, diamonds = outliers.

## Discussion

Our results show that *C. campestris* does not synchronize its flowering time with the host and that it can induce flowering in the absence of a host FT signal. We also found that *C. campestris*, as well as *C. australis*, encode and express endogenous *FT* and *FD-like* orthologs. The *CcFD-like* and *CaFD-like* genes are expressed in all tissues of the parasitic plant, similar to the *FD* homologs in rice and the *NtFD1* and *NtFD2* genes in tobacco (26, 32). In most species, *FT* genes are expressed primarily in the leaves, but also in other tissues (26–28, 32, 33). Accordingly, *FT* expression not only depends on endogenous cues such as hormones and plant age, and seasonal cues such as temperature or photoperiod, but also on carbon assimilation rates and, ultimately, sugar levels (34–36). The expression of *Cuscuta FT* homologs was only detected in the haustoria, not in the other tissues. In parasitic plants, sensing the physiological status of the host is likely to facilitate successful reproduction by ensuring a sufficient energy supply for growth, floral induction, seed maturation, and ripening. Indeed, during the *in vitro* cultivation of *C. campestris* excised stem tips only require sugars for growth and flowering (24, 25). Noteworthy, host-free grown cultures of *Cuscuta reflexa* respond to photoperiodic signals for floral induction (37, 38).

Floral transition genes have been studied in many species other than Arabidopsis, revealing numerous alternative genetic networks that regulate flowering time (20). In tobacco (Solanaceae), several FT homologs are produced, some of which act as floral activators and others as repressors (26–28). In potato (*Solanum tuberosum*), a close relative of tobacco, FT homologs not only control sexual reproduction but also vegetative reproduction, by coordinating the floral transition and tuberization (39). Sugar beet (*Beta vulgaris*; Chenopodiaceae) has no functional CO ortholog and its FLC homolog is not stably downregulated after vernalization, suggesting it does not play a major role in the vernalization response (40, 41). Instead, sugar beet produces a pair of FT homologs (the activator BvFT1 and the repressor BvFT2) that are differentially expressed in annuals and biennials (42). Their expression is controlled jointly by BOLTING TIME CONTROL1 (BTC1), encoding a pseudoresponse regulator protein, and DOUBLE B-BOX TYPE ZINC FINGER PROTEIN 19 (BBX19), which probably integrates temperature and photoperiod signals (43, 44). These are three cases, among many, in which floral regulating pathways diverge from the Arabidopsis model. The respective variations showcase adaptations that have evolved in the course of diversification among angiosperms colonizing different environmental niches (25).

An earlier claim that the *Cuscuta* spp. flowering time network is impaired was based on the inferred loss of genes with key roles in the regulation of *Arabidopsis* flowering time, such as CO and FLC (6, 22). However, CO does not have a conserved role as a regulator of photoperiod-dependent floral induction outside the Brassicaceae (Fig. S3), and the pathway evolved from CO-like (COL) genes following gene duplication within this family (45). Moreover, the vernalization response (mainly controlled by FLC in *Arabidopsis*) has convergently evolved multiple times after the continental drift (46). Besides, vernalization is not required for flowering of *C. australis*, so floral induction is unlikely to depend on a master regulator of the vernalization response.

*Cuscuta* plants perceive the physiological status of their host through their haustoria (47). These parasitic organs connect to the host xylem and phloem and provide a conduit through which *Cuscuta* acquires water, nutrients (such as sugars, amino acids and minerals), hormones, and a diverse set of host metabolites and macromolecules. These are transported in water and phloem sap across symplastic junctions at the parasite–host interface, where incoming signals induce the expression of response genes – for example, sucrose is known to promote flowering by activating *FT* genes (48). A combination of direct experimental and circumstantial evidence therefore suggests that the host provides a physiological cue to induce flowering in the parasite, but the genetic components of the parasite’s floral induction pathway are endogenous. Further studies are needed to identify the host-plant signals proposed by this hypothesis.

We found that the parasite’s FT-FD module is intact (e.g., Fig. 1, 2, 3, S10). CcFD-like1 and CaFD-like were shown to interact with CcFT1/CcFT2 and CaFT (Fig. S10), respectively, but also with NtFT5. The high degree of conservation of the FT-FD module across species boundaries has been demonstrated before (26, 30), including interspecific interactions between *Arabidopsis* (AtFD) and FT proteins in tobacco (NtFT1–4) and *Sorghum bicolor* (FT1, FT8 and FT10). None of these species form natural grafts with each another or exhibit any form of co-dependence.

Finally, we confirmed that CcFT1, CcFT2 and CaFT are functional floral activators by showing each was able to complement the non-flowering phenotype of our *Ntft4^−^Ntft5^−^* tobacco mutant lacking both endogenous floral activator FTs (Fig. 3, S11, S12). Although our data clearly show that *C. campestris* does not need host FT proteins for floral induction, the genetic and physiological factors controlling *Cuscuta* spp. *FT* and *FD* gene expression remain unclear. Based on our results and those reported previously (6), we hypothesize that *Cuscuta FT* gene expression, and thus flowering, depend on the nutrient supply or a phytohormonal signal provided by the host. The most likely physiological signal for FT activation is sugar, which the parasite obtains directly through the phloem sap. Sucrose and trehalose-6-phosphate both activate FT-dependent floral induction (48–50). Furthermore, the phloem connection and phloem sap perception occur in the haustoria, which may explain why endogenous *Cuscuta* FT is expressed primarily in this organ (Fig. 2; Table 1). There is also bioinformatic and experimental evidence for the retention of photoperiod-induced flowering in *Cuscuta* spp. These parasitic plants still produce sensors for both shortwave and long-wave light (Fig. S3) that act upstream of FT. This aligns with *in vitro* cultivation protocols, which use photoperiod shifts to induce flowering in host-free *Cuscuta* plants (37, 38).

Temperature shifts may be relevant for flower induction as well. Yet, either pathway would be regulated by proteins other than CO or FLC as they both diverge substantially in *Ipomoea* (Solanales), the closest non-parasitic relatives of *Cuscuta* (Fig. S3). However, these upstream regulators of FT are challenging to evaluate because SD conditions and cold temperatures also reduce the rate of photosynthesis and affect the physiological status of the host, including the rate of sugar transfer to the parasite. It is also difficult to disentangle signals that independently activate flowering in the parasite (e.g., hormones and the parasite age) from those influencing the flowering time of the host. This may also underly discordant reports of flowering time synchronization between *Cuscuta* spp. and its various hosts (51). It is therefore unclear whether the host contributes to floral induction in *Cuscuta* spp., and if so, how this is achieved and to what extent. We have demonstrated that *C. campestris* and *C. australis* possess an endogenous FT-FD module, which can serve as a reference point to help identify the upstream regulators and downstream targets.

## Materials and methods

### Plant material and cultivation conditions

This study used tobacco of the variety *N. tabacum* L. cv. Petit Havana SR1, *Ntft5^−^* mutant plants originating from the nullizygous T_2_ progeny of line #78 without the pDECAS9-*NtFT5*_ex I-147.169bp_ transgene (28), and *Glycine max* cv. Summer Shell, which were cultivated either under LD conditions in a climate-controlled greenhouse (16-h photoperiod, artificial light switched on if natural light fell below 700 μmol m^−2^ s^−1^, 22–25 °C under light, 19–25 °C in the dark) or under SD conditions in phytochambers (8 h photoperiod, 200 μmol m^−2^ s^−1^, 25–27 °C under light, 20 °C in the dark). *Cuscuta* spp. seeds, harvested from living collections of previous study material (21, 22), were sulfuric acid-treated before germination and infection of host plants. A low red/far-red light ratio, facilitating parasitism (52), was established using far-red (730-nm) LEDs with a photon flux density of 281.4 μmol m^−2^ s^−1^ in addition to the standard greenhouse lights until the first haustorial connection had developed.

### Generation of non-flowering Ntft4^−^Ntft5^−^ double knockout plants

The binary construct pDECAS9-*NtFT4*_ex I-59.79 bp_ was introduced into *Agrobacterium tumefaciens* by electroporation. The bacteria were then used for the stable transformation of *Ntft5^−^* plants (28). The presence of mutations in the *NtFT4* and *NtFT5* genes was verified by Sanger sequencing. For cloning and genome editing, we used recently described vector systems and experimental procedures (27, 53–58); SI Appendix, Materials and Methods.

### *Generation of Ntft4^−^Ntft5^−^ plants overexpressing* CcFT

The complete coding sequences of *CcFT1* and *CcFT2* were placed under the control of the strong and constitutive CaMV35S promoter in binary plasmids using the primers listed in Table S4. Strains of *A. tumefaciens* harboring these plasmids were used for the stable transformation of *Ntft4^−^Ntft5^−^* double knockout plants. Transgene integration was verified by PCR and transgene expression levels were determined by qRT-PCR.

### Bioinformatics and statistics

Based on a reference set of 295 protein-coding genes associated with flowering time regulation in *Arabidopsis thaliana* (23), we used blastX and exonerate to comparatively analyze gene conservation between selected eudicot species, *Cuscuta campestris*, and *C. australis*, the latter two partly reannotated using BRAKER (59) and syntenic block analysis (60). In addition, *Cuscuta* spp. transcriptomes (Table S1) were assembled *de novo* with the Trinity RNA seq pipeline (61), and FT, FD-like, as well as selected FT interacting genes were mapped using *bowtie2* (62). Gene trees for FT and FD-like sequences of *Cuscuta* spp. and other angiosperms (Tables S2 and S3) were reconstructed from *mafft* alignments (63) using RAxML (64). Furthermore, we examined the extended FT gene region using domain-based annotation of transposable elements (DANTE) with auto-filtering of the results. Data from genetic experiments were statistically analyzed with ANOVA, Tukey’s post hoc test, and Student’s t-tests using OriginPro2022 (details: SI Appendix, Materials and Methods).

## Data availability

All sequence data reported herein, including target-assembled protein-coding genes, alignment files, and gene family trees, as well as unscaled figures have been published in standardized, reusable *.fasta, .nexus*, and *.newick* file formats in the Dryad Data Repository, https://doi.org/10.5061/dryad.jsxksn0dr. All newly generated sequence data have been deposited at NCBI Genbank under accession numbers OP995431-OP995444.

## Acknowledgments

The authors would like to thank Dr. Kirsten Krause (UIT The Arctic University of Norway) and Dr. Jianqiang Wu (Kunming Institute of Botany, China) for providing plant material and seeds of *C. campestris* and *C. australis*, respectively. We thank Alina Griese for technical and horticultural assistance. We also thank Michael Lahme, Andreas Wagner and Jost Muth (Fraunhofer IME, Germany) for technical assistance. This study was made possible through financial support from the German Science Foundation (DFG, WI4507/3-1 to S.W.) and the Fraunhofer internal funding program PREPARE to G.A.N./D.P.

## Author contributions

S. Mäckelmann: Conceptualization; Investigation; Validation; Visualization. A. Känel: Conceptualization; Investigation; Validation; Visualization; Writing. L. M. Kösters: Investigation; Data analysis. P. Lyko: Investigation; Data analysis. D. Prüfer: Funding acquisition; Supervision; Writing-Reviewing & Editing. G. A. Noll: Conceptualization; Project administration; Writing-Reviewing & Editing. S. Wicke: Conceptualization; Data analysis, Funding acquisition; Project administration; Visualization; Writing-Reviewing & Editing.

