## Supplementary material for "Gene complementation analysis suggests that dodder plants (*Cuscuta* spp.) do not depend on the host FT protein for flowering": SI Appendix

**This PDF file includes:**

Supplementary text
Figures S1 to S13
Tables S1 to S4
Legends for Datasets S1 to S2
SI References

**Other supplementary materials for this manuscript include the following:**

Datasets S1 to S2

### Supplemental Materials and Methods

**Plant material and cultivation conditions**. All tobacco plants used in this study were the variety *N. tabacum* L. cv. Petit Havana SR1. The *Ntft5*⁻ plants were the nullizygous $T_2$ progeny of line #78, which no longer carries the pDECAS9-*NtFT5*$_{ex\ I\text{-}147.169bp}$ transgene (1). If not stated otherwise, tobacco or soybean (*Glycine max* cv. Summer Shell) seeds were sown and the plants were cultivated in standard soil under LD conditions in the greenhouse (16-h photoperiod, artificial light switched on if natural light fell below 700 µmol $m^{-2}$ $s^{-1}$, 22–25 °C under light, 19–25 °C in the dark), or under SD conditions in phytochambers (8-h photoperiod, 200 µmol $m^{-2}$ $s^{-1}$, 25–27 °C under light, 20 °C in the dark). For *Agrobacterium*-mediated transformation, wild-type tobacco plants, nullizygous *Ntft5*⁻ or nullizygous *Ntft4*⁻*Ntft5*⁻ plants were germinated and grown (LD, 16-h photoperiod, 100 µmol $m^{-2}$ $s^{-1}$, 23 °C) on sterile MS medium. *Cuscuta* spp. seeds, obtained from living collections at WWU Münster and the Späth-Arboretum Berlin, originating from material of UIT Arctic University of Norway and the Kunming Institute of Botany (2, 3), were incubated in 97% sulfuric acid for 20 min and then washed with tap water 3–4 times. Afterwards, the seeds were placed on wet filter paper in Petri dishes and transferred to the greenhouse for germination. Seedlings (2–4 cm in length) were placed near the stems of tobacco at the 2–3 true leaf stage, whereas soybean was parasitized as soon as the plants reached ~10 cm in height. A low red/far-red light ratio, facilitating parasitism (4), was established using far-red (730-nm) LEDs with a photon flux density of 281.4 µmol $m^{-2}$ $s^{-1}$ in addition to the standard greenhouse lights until the first haustorial connection had developed.

**Identification and validation of genomic and coding sequences.** Genomic DNA was isolated from *C. campestris* using the NucleoSpin Plant II Kit (Machery Nagel, Düren, Germany) and the genes were amplified in overlapping fragments using Phusion High-fidelity polymerase (Thermo Fisher Scientific) and the primers listed in Table S3. Gene fragments were subcloned into pCRIITopo (Thermo Fisher Scientific) and sequenced. To identify the 3′-UTR by 3′-RACE, RNA was isolated from *C. campestris* or *C. australis* haustoria using the PureLink RNA Mini Kit (Thermo Fisher Scientific) and reverse transcribed using Superscript II reverse transcriptase (Thermo Fisher Scientific) and the primers listed in Table S3. The cDNAs were used as templates for PCR using Phusion High-fidelity polymerase and the primers listed in Table S4. Based on the identified 3′-UTR sequence, *CcFT1*, *CcFT2* and *CaFT* full-length coding sequences including parts of the 3′-UTR were amplified and verified by sequencing.

**Quantitative real-time PCR.** Plant tissues were harvested, snap-frozen in liquid nitrogen and ground using either an MM400 bead mill (Retsch, Haan, Germany) or with a mortar and pestle under liquid nitrogen. Tobacco RNA was extracted using the innuPREP Plant RNA kit (Analytik Jena, Jena, Germany) and *Cuscuta* spp. RNA was extracted using the PureLink RNA Mini Kit. Residual genomic DNA was digested using the TURBO DNA-free kit (Thermo Fisher Scientific). Following reverse transcription using PrimeScript RT master mix (Takara Bio), gene expression was analyzed by quantitative real-time PCR using Kapa SYBR Fast qPCR Master Mix (Merck, Darmstadt, Germany), the primers listed in Table S3, and the CFX96 Real-Time System (Bio-Rad Laboratories, Hercules, CA, USA). Each reaction was carried out in technical triplicates. Specificity was ensured by melt curve analysis, the sequencing of PCR products, and by including no-template and no-reverse-transcription controls. Individual PCR efficiency was determined using LinReg PCR v2017.0 and relative gene expression levels were normalized to *EF1α* and *Actin* (*Cuscuta* spp.), *EF1α* (*N. tabacum*), or *F-Box* (*G. max*).

**Cloning.** For the heterologous expression of *CcFT* genes, we amplified the CaMV35S promoter sequence including the TMV Ω leader from pFGC5941-GW (www.chromdb.org) using the splicing by overlap extension (SOE)-PCR technique to eliminate the internal XhoI site. The amplicon was then transferred to the SmaI/XhoI sites of pLab12.10 (5) or pLab12.1 (6) to generate pLab12.10Q35S-P and pLab12.1Q35S-P, respectively. The vector backbones carry different resistance cassettes (pLab12.10 confers kanamycin resistance and pLab12.1 confers phosphinothricin resistance). *CcFT1* and *CcFT2* were amplified from cDNA using primers with attached restriction sites (Table S3) and were transferred to pLab12.10Q35S-P and pLab12.1Q35S-P, respectively, by restriction and ligation. For the targeted editing of *NtFT4*, the binary construct pDECAS9-*NtFT4*$_{ex\ I\text{-}59.79\ bp}$ was cloned as previously described
(1). The original plasmids were kindly provided by Holger Puchta (Karlsruhe Institute of Technology, Karlsruhe, Germany). For BiFC assays, *CcFT1*, *CcFT2* and *CcFD-like1* were amplified from cDNA using primers with attached restriction sites (Table S3) and were transferred to pENTR4 (Thermo Fisher Scientific) by restriction and ligation. Subsequent transfer to pBatTL vectors was achieved by Gateway recombination (pBatTL plasmids were

kindly provided by Joachim Uhrig and Guido Jach, University of Cologne, Cologne, Germany). BiFC constructs containing *NtFD1* and *NtFT5* were available from previous studies (7).

***Agrobacterium*-mediated stable transformation of tobacco.** Tobacco transformation was carried out using the leaf disc method with *Agrobacterium tumefaciens* strain LBA4404 for heterologous gene expression, or EHA105 for CRISPR/Cas9-mediated genome editing. The appropriate binary vectors were introduced into these strains by electroporation. For the selection of transgenic plants, MS medium was supplemented with 100 mg/L kanamycin or 3 mg/L phosphinothricin, as appropriate. After callus regeneration and rooting in sterile culture medium, independent transgenic plant lines were cultivated in soil under LD conditions in the greenhouse as described in the main article.

**Identification and screening of genome-edited plants.** The *NtFT5* locus in the mutant plants (*Ntft5*$^{-}$ and *Ntft4*$^{-}$ *Ntft5*$^{-}$) was analyzed by isolating genomic DNA using the NucleoSpin Plant II Kit and amplifying exon I using Phusion High-fidelity polymerase and the primers listed in Table S3. The amplicon was verified by sequencing. After regenerating *Ntft5*$^{-}$ plants transformed with pDECAS9-*NtFT4*$_{\text{ex I-59.79 bp}}$, transgene integration was verified by the isolation of genomic DNA followed by PCR using the 2x *i*-MAX Mastermix (INtRON Biotechnology, Seongnam, Korea) and the primers listed in Table S3. For the analysis of genome editing in $T_0$ plants, parts of exon I were amplified using Phusion High-fidelity polymerase and the primers listed in Table S3. The amplicons were transferred to pCRIITopo and sequenced. Promising plants were further cultivated for seed production and the $T_1$ progeny was analyzed again regarding knockout mutations. For the analysis of genome editing in $T_1$ plants, parts of exon I were amplified using the Phusion High fidelity polymerase and the primers listed in Table S3. The amplicons were either transferred to pCRIITopo for sequencing (if the plants carried different mutations in their two *NtFT4* alleles) or were sequenced directly (if the same mutation was present in both alleles).

**Phenotyping.** For tobacco, we counted the number of days from seed sowing to full opening of the first floral bud, and the number of leaves when the first floral bud fully opened. For *Cuscuta* spp., we counted the number of days from the formation of the first haustorial connection until the first floral bud had fully opened.

**Infiltration of *N. benthamiana*.** For the transient expression of BiFC constructs, *A. tumefaciens* strain GV3101 pMP90 was transformed with the corresponding binary vectors by electroporation. *N. benthamiana* plants were cultivated in the greenhouse (16-h photoperiod) until they were 4–5 weeks old. The leaves were then infiltrated with *A. tumefaciens* strain GV3101 pMP90 carrying the appropriate plasmids ($OD_{600}$ = 0.25) and *A. tumefaciens* strain C58C1 ($OD_{600}$ = 0.3) carrying the pCH32 helper plasmid and the pBin61 plasmid encoding the RNA silencing suppressor p19 from tomato bushy stunt virus (8). Plants were cultivated under continuous light for 3–4 days, and leaf discs were screened for fluorescent cells in the abaxial epidermis.

**Microscopy.** For BiFC experiments, fluorescence was analyzed using a STELLARIS 8 confocal laser scanning microscope (Leica Microsystems, Wetzlar, Germany) at excitation/emission wavelengths of 549/569–629 nm for reconstituted mRFP. Interaction was confirmed if at least five independent images showing fluorescence were captured.

**Bioinformatic recovery of flowering time genes.** We extracted the full set of potentially functional flowering genes and corroborate gene losses in *Cuscuta campestris* and *C. australis* relative from the genomic sequences of both species from NBCI Genbank (Bioprojects: PRJEB19879; PRJNA394036), in addition to assembling and querying RNA seq covering key developmental stages (Table S1). Genome data alongside their published annotations of 13 closely related eudicot species (*I. triloba* – PRJNA574454; *Beta vulgaris* – assembly EL10_1.0; *Spinacia oleraceae* – assembly ASM200726v1; *Nicotiana tabacum* – assembly Ntab-TN90; *Nicotiana sylvestris* – assembly Nsyl; *Nicotiana tomentosiformis* – assembly Ntom_v01; *Vaccinium darrowii* – assembly USDA_Vadar_1.0_pri; *Solanum lycopersicum* – assembly SL3.1; *Solanum tuberosum* – assembly SolTub_3.0; *Erythranthe guttata* – assembly Mimgu1_0; *Olea europaea var. sylvestris* – assembly O_europaea_v1; *Lactuca sativa* – assembly Lsat_Salinas_v8; *Daucus carota subsp. sativus* – assembly ASM162521v1) were included for comparative-evolutionary analyses of gene conservation. As genomic data were of significantly different quality regarding annotation version, we used a two-stage flowering time gene analysis, in which a primary query for flowering time genes in all species was conducted on the raw genomic assembly (annotation-free) and hits were then cross-validated and refined using readily available gene annotations/predictions. As both *Cuscuta* spp. differ significantly in their coding capacity (2, 3, 9), we also reciprocally searched their genomes for missing protein-coding genes using blastX (e-value cutoff: $10^{-3}$) and exonerate (protein2genome mode; min length coverage: 70 %), and compared these data to the closest, nonparasitic relative *Ipomoea triloba*, which belongs to the same family as *Cuscuta* spp. (Convolvulaceae). As this approach retrieved many originally "missing" genes (data published in the DRYAD Data repository, https://doi.org/10.5061/dryad.jsxksn0dr), we used the BRAKER pipeline to reannotate *Cuscuta* spp. and *Ipomoea nil* (assembly Asagao_1.1; Bioprojects: PRJNA344313, PRJDB4356) for a direct comparison of the new *ab initio* gene model predictions; training of Augustus of the BRAKER pipeline followed the steps described for *C. campestris* (2), whereby we also used the odp10 protein database plus *I. nil* and *I. triloba* proteins. Our annotation revealed an almost perfect 2:1 gene annotation ratio between *C. campestris* and *C. australis*. Since an analysis of syntenic blocks across *Cuscuta australis*, *C. campestris*, and *Ipomoea triloba* computed using the Doerr & Moret (2017) pipeline (10) found counterparts with conserved genomic orientation on all genomic fragments longer than three million bases in both parasite species (DRYAD Data repository, https://doi.org/10.5061/dryad.jsxksn0dr), we conclude from these data that the genome assemblies may be available in collapsed vs. uncollapsed allelic/pseudochromosomal configuration, rather than reflecting a ploidization event since the parasite species' evolutionary split. For the purpose of screening for flowering genes, we merged these new gene sets with the corresponding original ones and removed redundancy in cases of perfect matches. The genomic sets were used to seed-search for 295 protein coding genes implicated in flowering time regulation using the *exonerate* program in protein2genome mode (https://www.ebi.ac.uk/about/vertebrate-genomics/soft-ware/exonerate), allowing for introns between 15 bp and 35 kb in length. Reference flowering time proteins were obtained from the FLOR-ID database http://www.phytosystems.ulg.ac.be/florid/ (11) and the Araport11 genome release of *Arabidopsis thaliana* (https://www.arabidopsis.org/index.jsp). We excluded all of the eleven miRNAs involved in flowering time regulation. In addition, the exonerate-based search for flowering time genes using the Arabidopsis reference gene set was repeated for transcriptome assemblies of *C. australis* and *C. campestris*, using all RNAseq data sets summarized in Table S1. *Cuscuta* spp. transcriptomes were assembled *de novo* with the Trinity RNA seq pipeline using default settings (12). Recovered hit sequence seeds of queried genes of *Cuscuta* spp. were then used to extend the region bidirectionally by iteratively mapping genomic and transcriptomic sequence data using *bowtie*2, whereby up to 10 iterations were performed. The exonerate wrapper and all retrieved seed sequence alignments have been deposited in the DRYAD Data repository, https://doi.org/10.5061/dryad.jsxksn0dr. To compare the divergence in sequence conservation of *Cuscuta* spp. flowering time genes with the corresponding homologous gene models of 13 other eudicot species, we ran *exonerate* in protein2genome mode (as above). Matched protein-coding gene regions were extracted, aligned using *mafft* in auto mode (13), and evaluated regarding the highest-scoring coding sequence length and sequence divergence, relative to functionally validated references of *Arabidopsis thaliana* (% conservation of gene models). Finally, we used phylogeny-aware domain-based examination, for which we extracted the full gene models matching reference FD and FT sequences from the combined genomic annotations. We build FT and FD alignments using *mafft* in the e-ins-i mode with a gap penalty of 1.93. From these data, we reconstructed the protein relationships with RAxML (14) with 200 bootstrap replicates, employing the WAG model of amino acid substitution with a gamma rate distribution.

**Transposable element analysis.** To check whether transposable element insertion might have rendered FT or FD nonfunctional, we used domain-based annotation of transposable elements (DANTE; https://github.com/kavonrtep/dante). To this end, we extracted the extended gene regions (-20 kbp and + 20kbp of the entire genes of both species), thus allowing us to retrieve full-length mobile DNA elements, if any. DANTE was run with the Viridiplantae 3.0 as taxon and protein database and a BLOSUM80 scoring matrix. Single hits were retained if they reached a similarity hit with database records of 80% or higher. DANTE's built-in *Protein Domain Filter* tool was then used on the full DANTE annotation and only domains with at least 35 % protein sequence identity between input and mapped protein from the database, 45 % minimum similarity, and 80 % minimum alignment length. Hits with more than three interruptions such as frameshifts and/or stop codons and insertions resulting in a domain length extensions of 1.2 between the new and the reference domain from the database were filtered. The resulting filtered and unfiltered evidence for transposable elements were extracted from GFF3 files.

**RNAseq based gene expression analysis.** For a RNAseq-based gene expression analysis, we extracted the full coding sequence models of our candidate FD and FT sequences, and, for comparison, those of the FT interacting genes SOC1, CO, LFY, TFL1, and GI from the combined genomic annotations of both *Cuscuta* species. We assessed gene expression of FT, FD, and our selected FT(-FD) interacting proteins by mapping all available RNA Seq data (Table S1) using *bowtie2* in local mode (15). The unsorted read alignments were parsed to calculate per-reference base coverage using the *pileup* function of the BBMap program suite (16; https://github.com/BioInfoTools/BBMap). As RNAseq based expression levels were extremely low for FT (Table 1), its mapping results were double-checked and confirmed by qRT-PCR using highly discriminating primers (see above; Table S3).

**Statistics.** All boxplots in the figures were prepared in OriginPro2022 (OriginLab Corporation, Northampton, MA, USA) using the default settings (center line = median; square = mean, box limits = upper and lower quartiles; whiskers= 1.5× interquartile range; diamonds = outliers). Statistical analysis was carried out using OriginPro2022. Equality of variances was determined by one-way analysis of variance (ANOVA), and pairwise comparisons were assessed using Tukey's *post hoc* test for multiple comparisons and Student's *t*-test for single pairwise comparisons, unless stated otherwise.

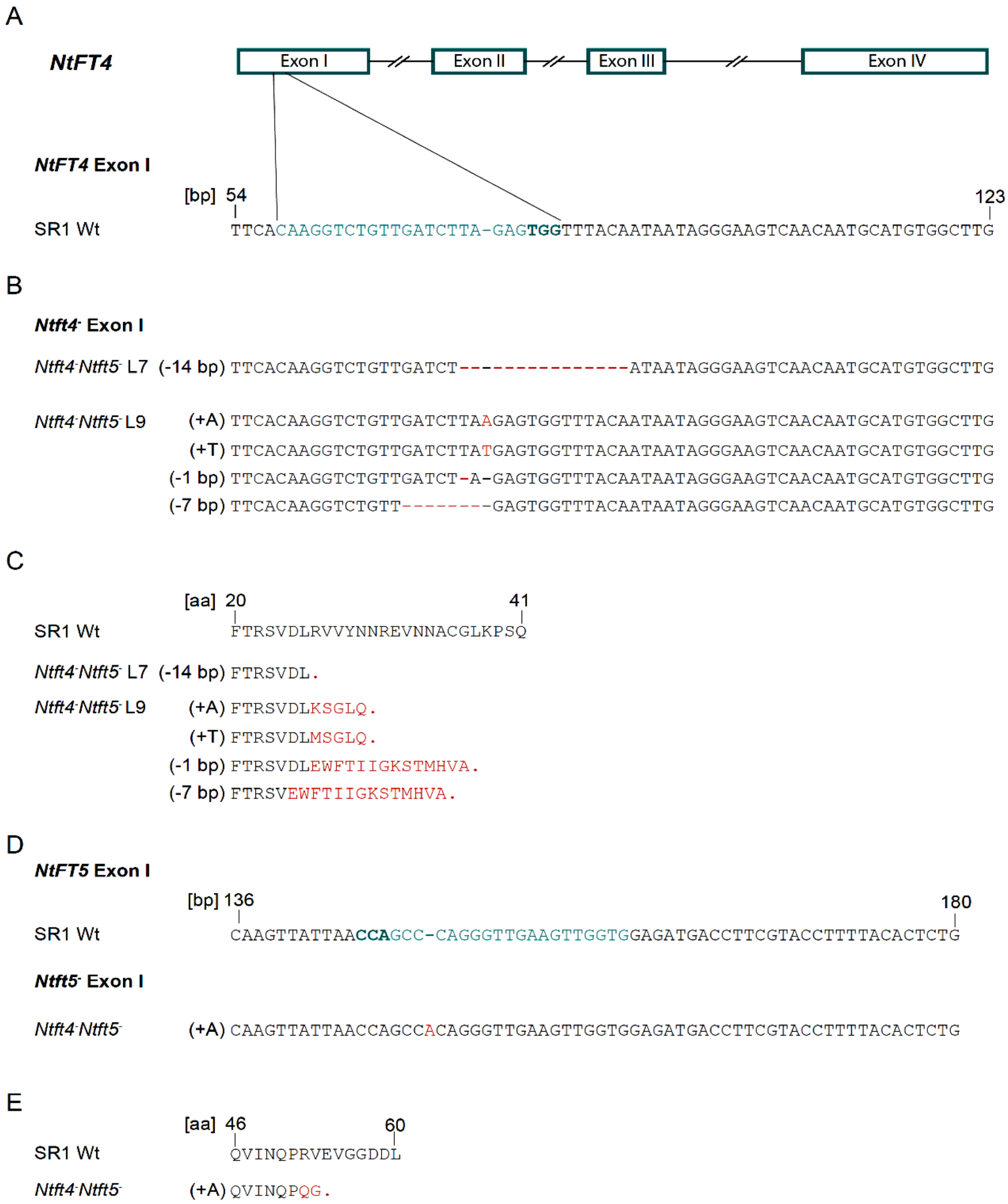


**Fig. S1. CRISPR/Cas9 genome editing of *Ntft5*⁻ plants by transformation with pDECAS9-*NtFT4*$_{\text{ex I-59.79 bp}}$.** **A)** Location of the *NtFT4*-specific protospacer (teal) and the protospacer adjacent motif (PAM; bold) in the 59–82 bp region of exon I in SR1 wild-type plants (SR1 Wt). Exons are shown as boxes, introns as lines. **B)** PCR-based screening of two independent transgenic lines (L7, L9) for mutated allelic variants of *NtFT4*. Amplicons were subcloned and sequenced. Insertions and deletions are highlighted in red letters. L7 progeny plants possess a homozygous 14-bp deletion, whereas in L9 progeny plants the two *NtFT4* alleles each harbor one of four different mutations. **C)** Sequence alignment of native NtFT4 and the protein variants encoded by the mutated alleles. Amino acid substitutions and premature stop codons are highlighted in red letters. **D)** Knockout of *NtFT5* by a single base pair insertion at position 153 in exon I, leading to a frameshift and a premature stop codon, as validated by PCR and direct sequencing of the PCR product. **E)** Sequence alignment of native NtFT5 and the protein variant encoded by the mutated allele. Amino acid substitutions and premature stop codon are highlighted in red letters.

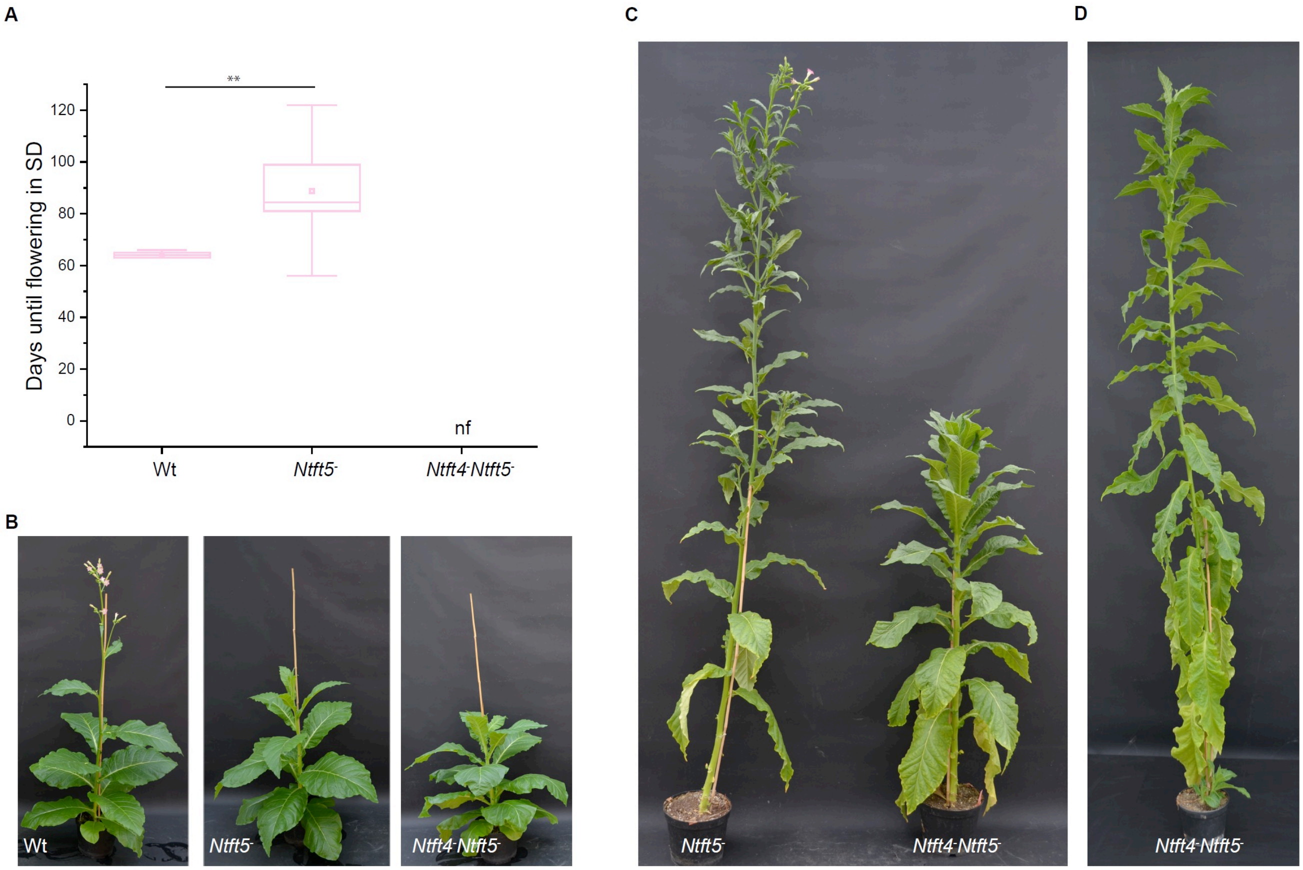


**Fig. S2. Phenotypic analysis of *Ntft4*$^{-}$*Ntft5*$^{-}$ plants. A)** Flowering time of SR1 wild-type plants (Wt), *Ntft5*$^{-}$ plants and *Ntft4*$^{-}$*Ntft5*$^{-}$ plants. Under short-day (SD) conditions, SR1 wild-type plants flowered on average after 64 days, whereas *Ntft5*$^{-}$ plants, which carry homozygous knockout mutations in *NtFT5* but still express *NtFT4* under SD conditions, flowered on average after 89 days. *Ntft4*$^{-}$*Ntft5*$^{-}$ plants, lacking both floral activators, did not flower at all under SD conditions up to the end of the experiment (122 days after sowing). Days until flowering were determined when the first bud had fully opened. ** Statistically significant difference as determined using Student's *t*-test with Welch's correction, $p < 0.01$ ($n = 10$). In the boxplots, center line = median, square = mean, box limits = upper and lower quartiles, and whiskers = 1.5× interquartile range. **B)** Phenotypes of Wt, *Ntft5*$^{-}$, and *Ntft4*$^{-}$*Ntft5*$^{-}$ plants cultivated under SD conditions 65 days after sowing. **C)** Phenotypes of *Ntft5*$^{-}$ and *Ntft4*$^{-}$*Ntft5*$^{-}$ plants cultivated under SD conditions 120 days after sowing. **D)** Phenotype of *Ntft4*$^{-}$*Ntft5*$^{-}$ plant cultivated under LD conditions 139 days after sowing.

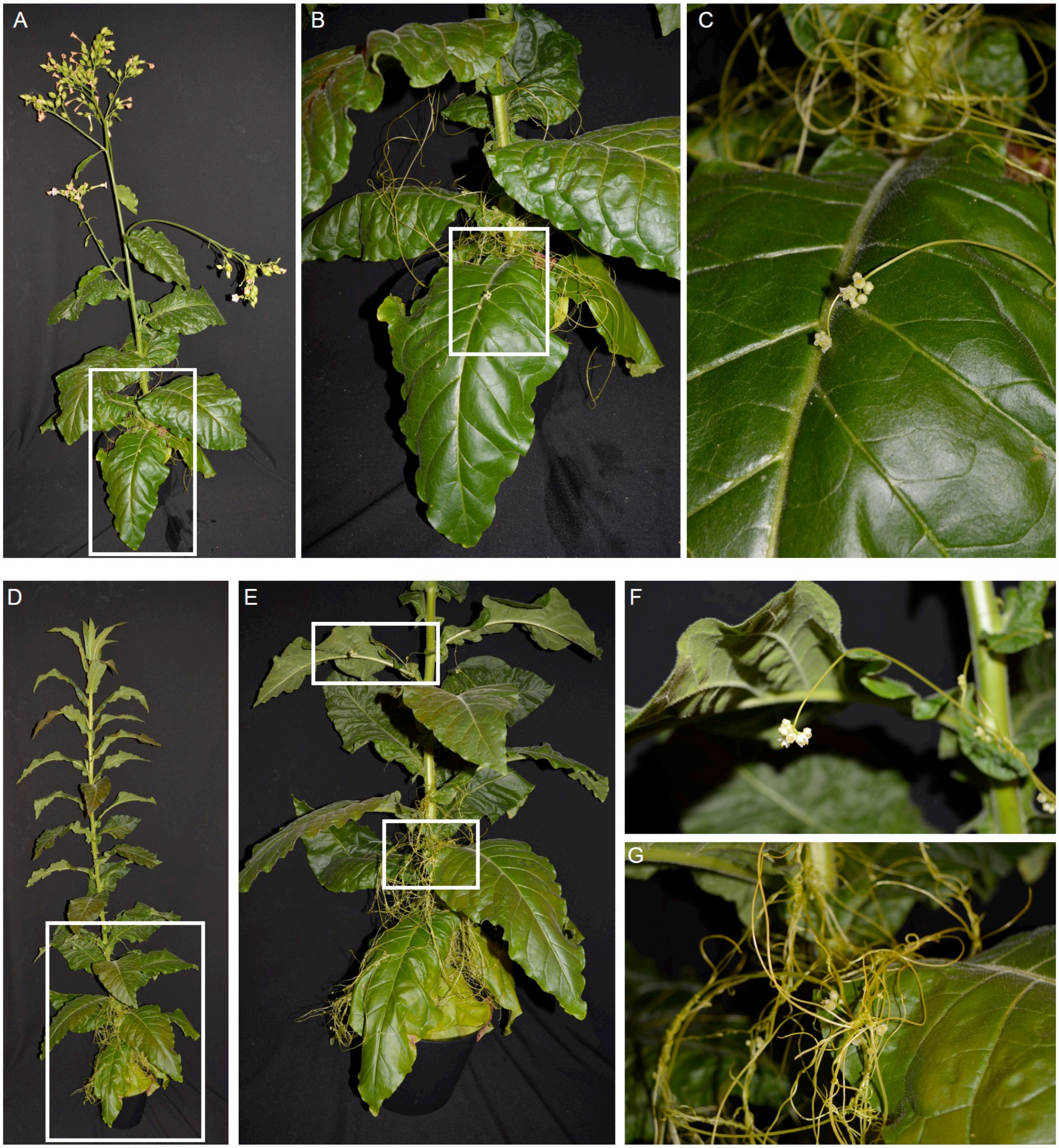


**Fig. S3. Flowering phenotypes of *C. campestris* parasitizing SR1 wild-type or non-flowering *Ntft4*$^{-}$*Ntft5*$^{-}$ plants.** *C. campestris* flowered similarly when parasitizing **(A-C)** SR1 wild-type or **(D-G)** *Ntft4*$^{-}$*Ntft5*$^{-}$ plants. Magnified areas from A (B, C) and D (E, F, G) are indicated by white squares. Photographs were taken 83 days after sowing.

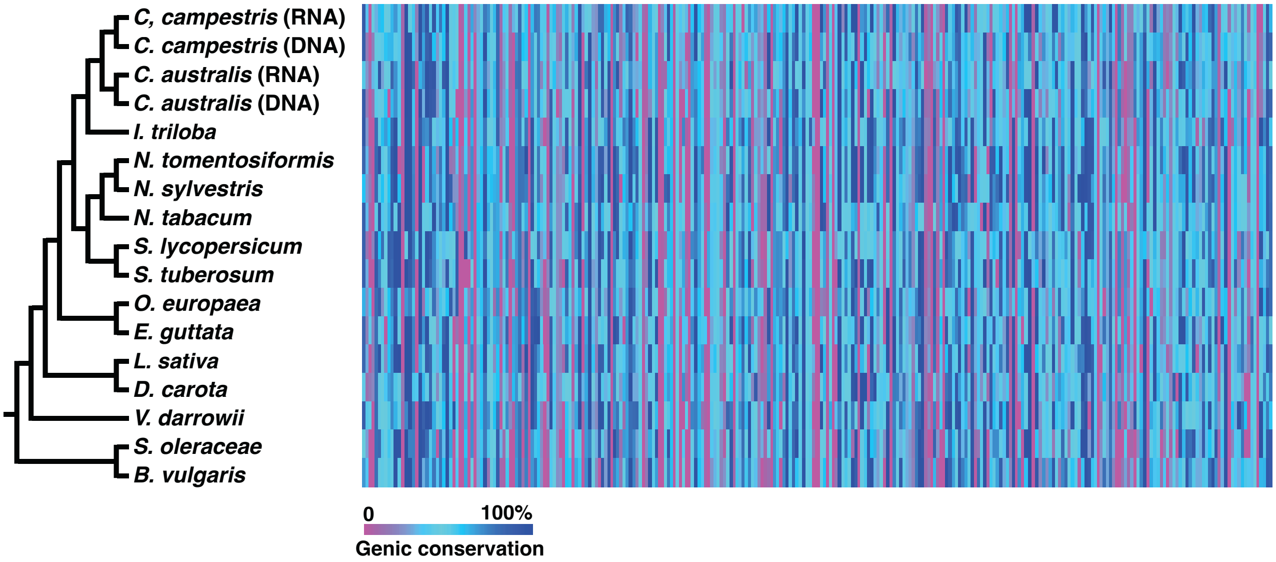


**Fig. S4. Graphical summary of genic conservation of flowering time genes in eudicots.** A heatmap shows the gene and reading frame conservation of flowering time genes in the genomes of *Cuscuta* spp. and 13 eudicots. Small blocks in columns represents one of 295 flowering time-associated protein coding genes. Blue indicates high similarity in length and translated amino acid sequence, whereas cyan and magenta colored blocks highlight genic drift or absence of a coding region, respectively. For *Cuscuta* spp., both genome (DNA) and transcriptome (RNA) assemblies were investigated, indicating that, in a few cases, gene models may be divergent and complex to predict, but presence of a flowering genes is likely on the mRNA level. Phylogenetic relationships are drawn according to those accepted by the Angiosperm Phylogeny Group (APG IV, 2016). Supplemental dataset 1 provides a detailed, gene-based overview. An unscaled version of this figures as well as the underlying species-specific gene fragment data are published in the DRYAD Data Repository, https://doi.org/10.5061/dryad.jsxksn0dr.

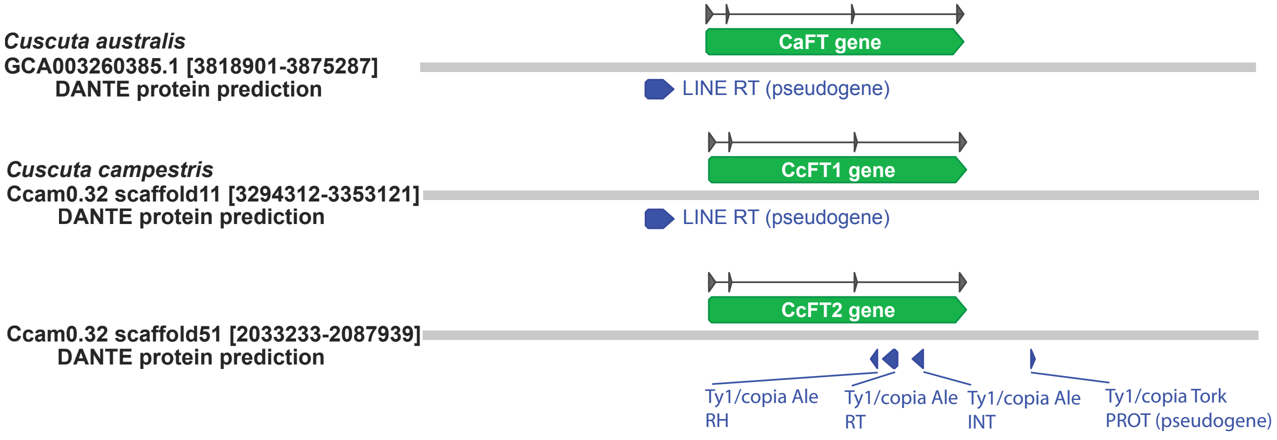


**Fig. S5. Graphical summary of FT gene regions.** The extended FT regions of *Cuscuta campestris* (CcFT1, CcFT2) and *C. australis* (CaFT) were examined for the presence of transposable elements shows a pseudogene of a LINE-type reverse transcriptase (RT) upstream of CaFT and CcFT1, whose coding sequence are identical. The third intron of CcFT2 carries a partial long-terminal repeat (LTR) retrotransposon of the the Ty1/copia Ale family, with partly intact genes for the ribonuclease (RH), reverse transcriptase (RT), and intergrase (INT). The group-specific antigen (GAG) and protease (PR) required for retrotranspososon activity and functionality are absent. A PR pseudogene with similarity to the Ty1/copia Tork family resides upstream of the fourth CcFT2 exon. The full list of filtered and unfiltered transposable element sequences, including protein translations of best hit fragments of TE domains are provided in Supplemental Dataset 2 and in the DRYAD Data Repository, https://doi.org/10.5061/dryad.jsxksn0dr.

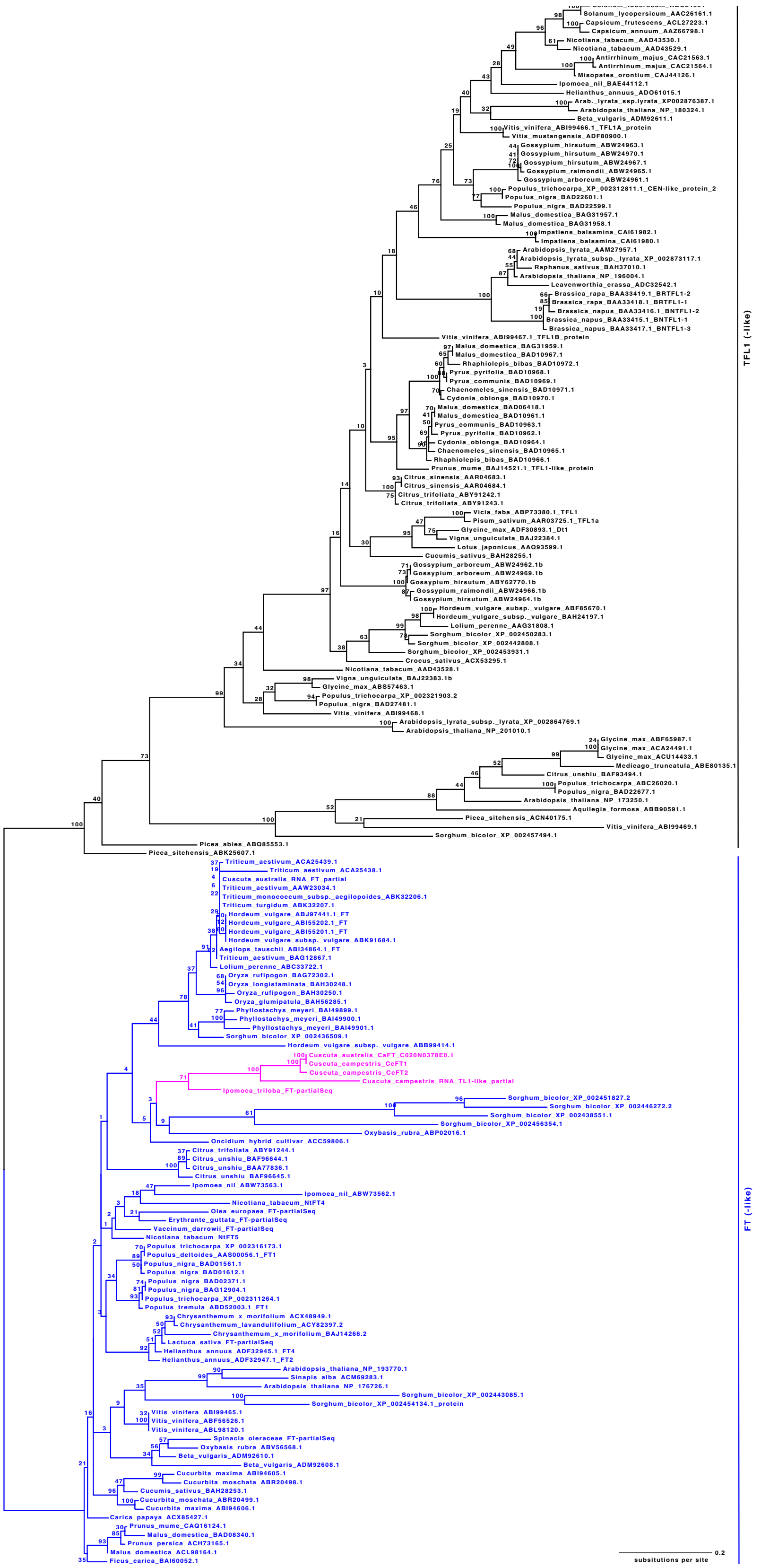


**Fig. S6. PEBP gene family tree.** Phylogenetic analysis shows that CcFT1, CcFT2, and CaFT (magenta-colored) cluster with validated FT homologs. Branch values indicate node support from 200 bootstrap replicates; scale bar marks substitutions per site. This tree is published in reusable Newick format in the DRYAD Data Repository, https://doi.org/10.5061/dryad.jsxksn0dr.

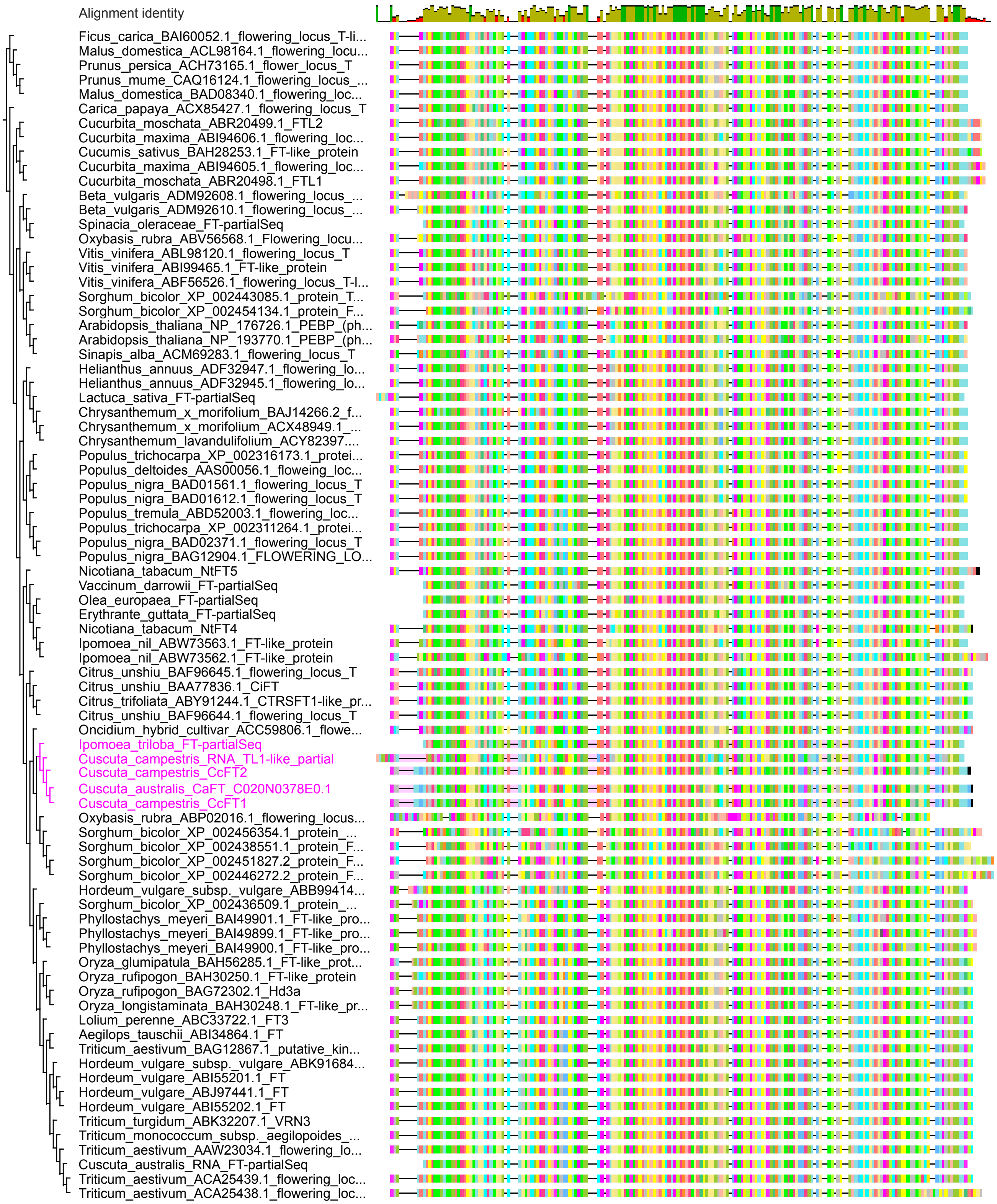


**Fig. S7. FT gene region.** Visualization of aligned sequences belonging to the FT-clade of the PEBP gene family, highlighting the conservation of the FT coding sequences from the *Cuscuta* spp. and *Ipomoea triloba* (magenta-colored). Full alignment is published in reusable Nexus format in the DRYAD Data Repository, https://doi.org/10.5061/dryad.jsxksn0dr.

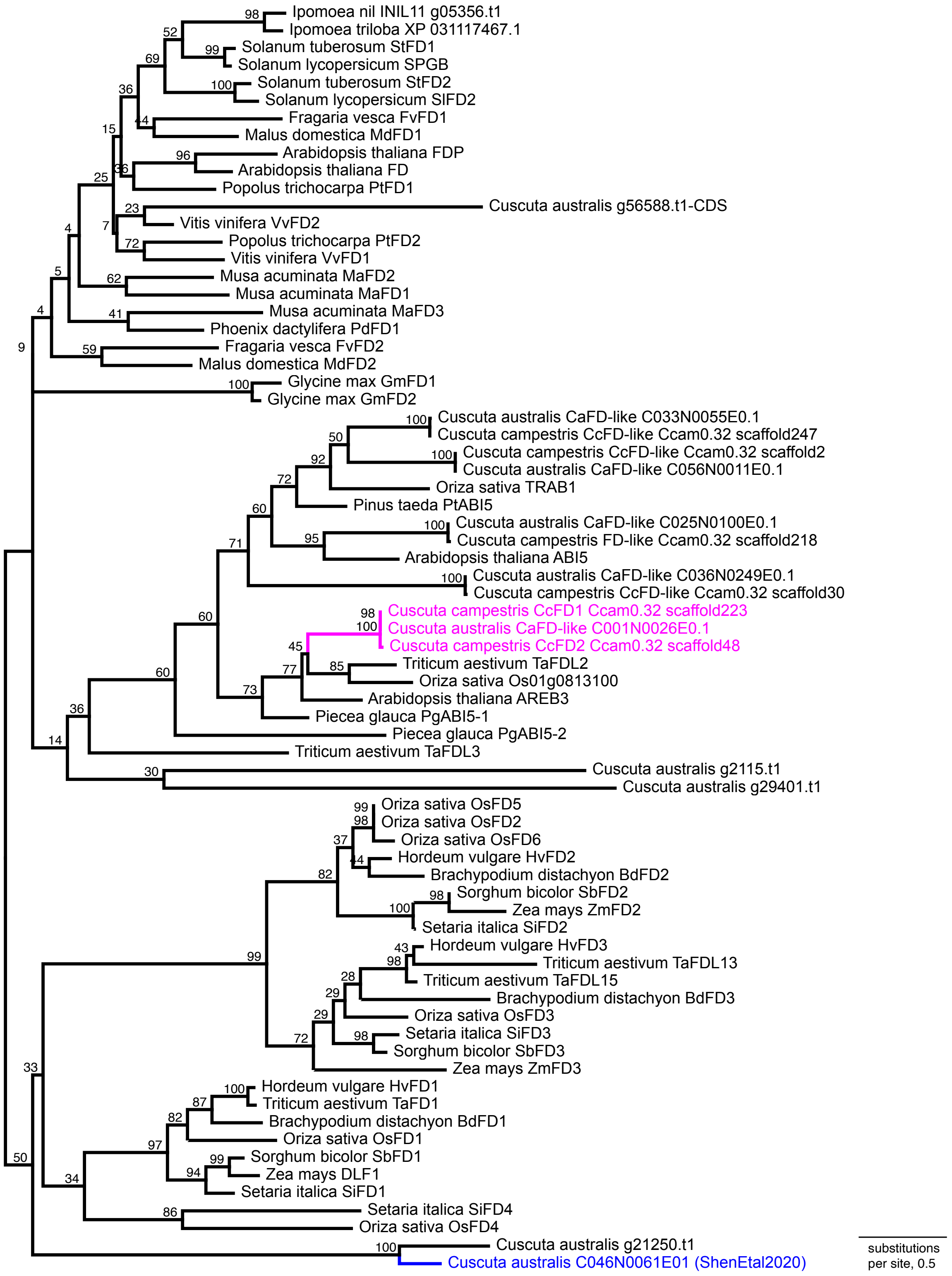


**Fig. S8. Gene tree of FD-like sequences.** Phylogenetic analysis shows the affinity of FD-like sequences to validated FD, AREB3, and ABI5 proteins within the bzip-family. The newly identified and tested FD-like genes are highlighted in magenta-colored; blue indicates a previously investigated FD-like gene of C. australis that was reported to not interact with endogenous *Cuscuta* spp. FT proteins (17). Branch values indicate node support from 200 bootstrap replicates; scale bar marks substitutions per site. This tree format is published in reusable Nexus and Newick format in the DRYAD Data Repository, https://doi.org/10.5061/dryad.jsxksn0dr.

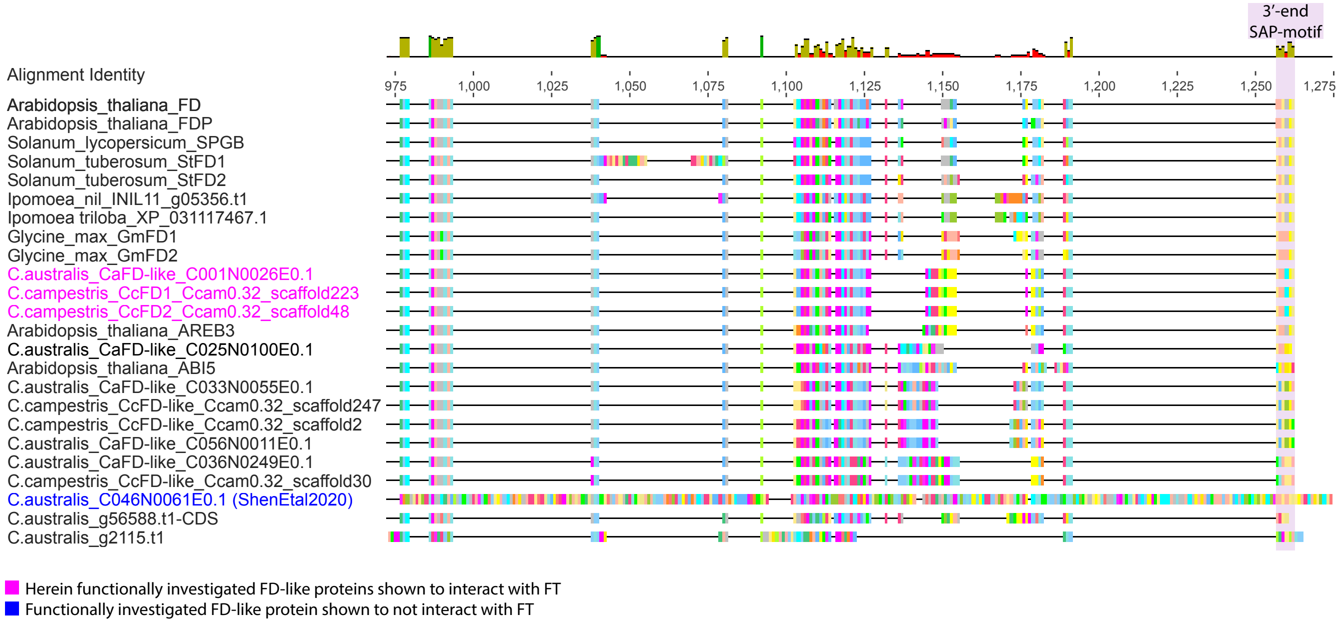


**Fig. S9. SAP-motif region of FD-like proteins.** Visualization of aligned sequences at the 3'-end of FD-like sequencing, showing the presence of a SAP-like motif in the newly identified and functionally tested CcFD1, CcFD2 and CaFD-like of *Cuscuta* spp. (magenta-colored), for which we showed an interaction with endogenous *Cuscuta* spp. FT proteins. For comparison, functionally validated FD and FD-related proteins from *Arabidopsis thaliana* and other angiosperms are included. Highlighted in blue is the sequence of the FD protein that was found to not interact with the endogenous FT of *Cuscuta* spp. previously (17). The complete alignment is published in reusable Nexus format in the DRYAD Data Repository, https://doi.org/10.5061/dryad.jsxksn0dr.

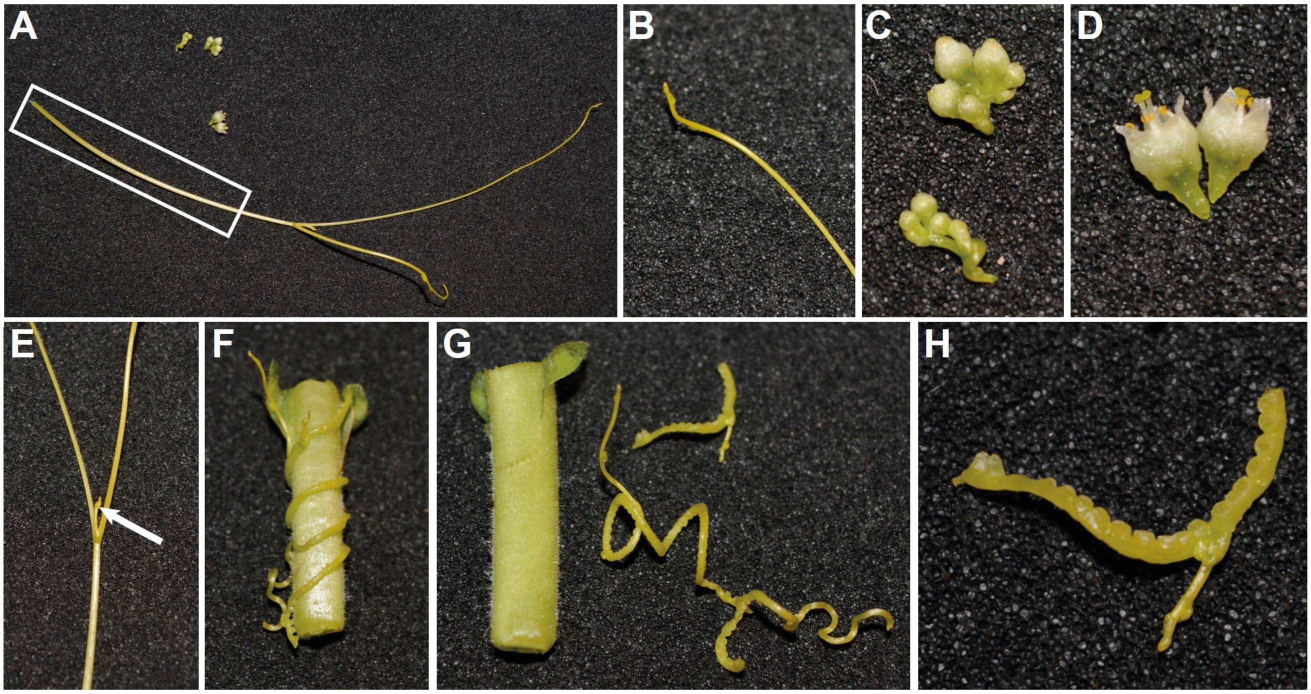


**Fig. 10.** ***Cuscuta*** **tissues harvested for qRT-PCR analysis. A)** For stem tissue (white square), we harvested ~8 cm of the twines. **B)** The stem tips include 2–3 cm of the apical stem region. **C)** Buds were harvested before they opened. **D)** Flowers were harvested when they were fully open. **E)** Branches were harvested with 2–3 cm of stem tissue in all directions, a scale-like leaf is indicated by an arrow. **F)** A haustorium connected to the host stem was carefully detached **(G)** and detached haustoria **(H)** were used for qRT-PCR analysis.

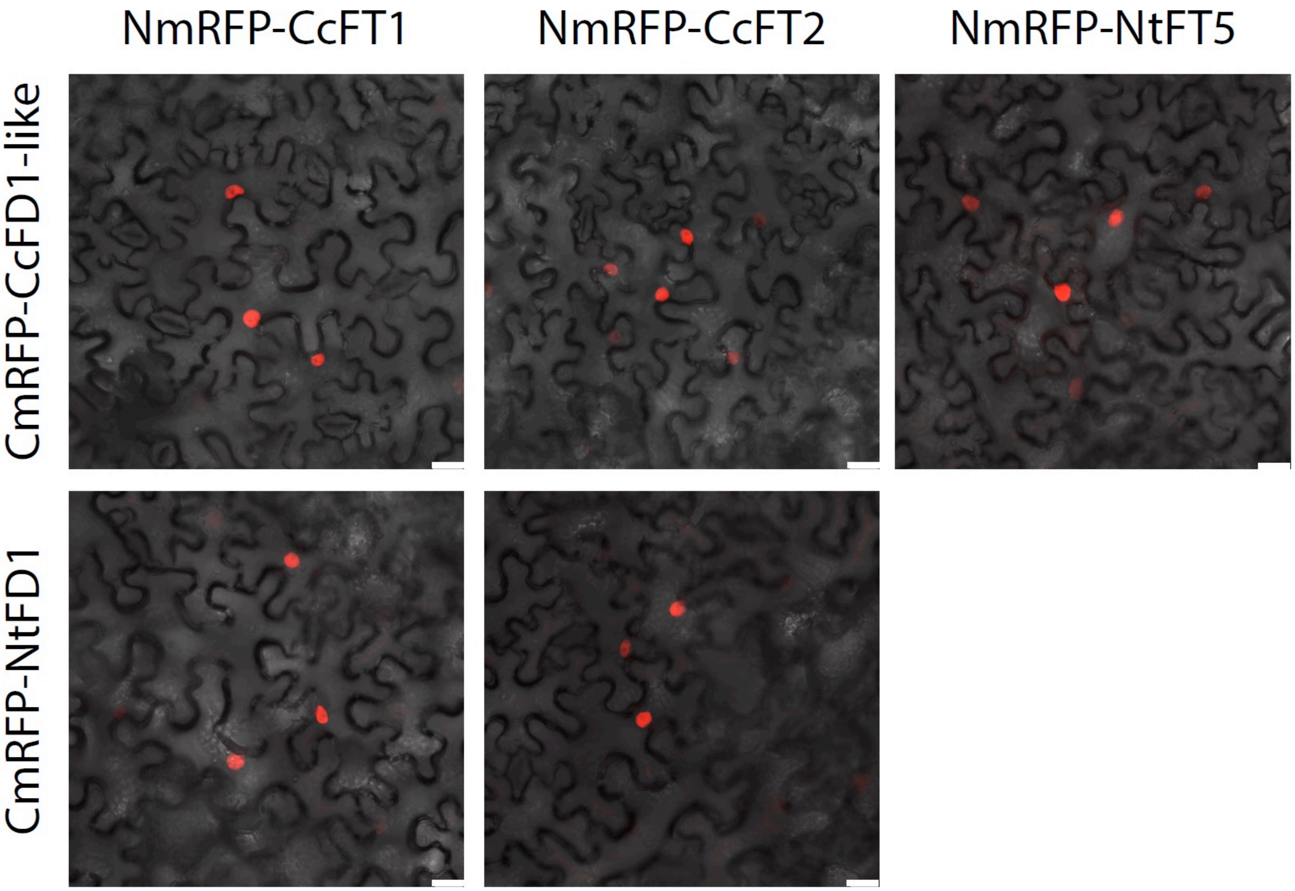


**Fig. 11. Bimolecular fluorescence complementation assay in *N. benthamiana* epidermal cells with *Cuscuta* and tobacco FD/FT homologs.** The FD homologs were C-terminally fused to the C-terminal mRFP1 fragment and the FT homologs were C-terminally fused to the N-terminal mRFP1 fragment. Interaction was confirmed by reconstituted mRFP fluorescence visible in the nuclei. N-terminal or C-terminal mRFP1 fragments served as negative controls, and were co-expressed with the corresponding FD or FT fusion proteins. We did not observe any nonspecific interactions in any combination. Scale bar = 20 µm.

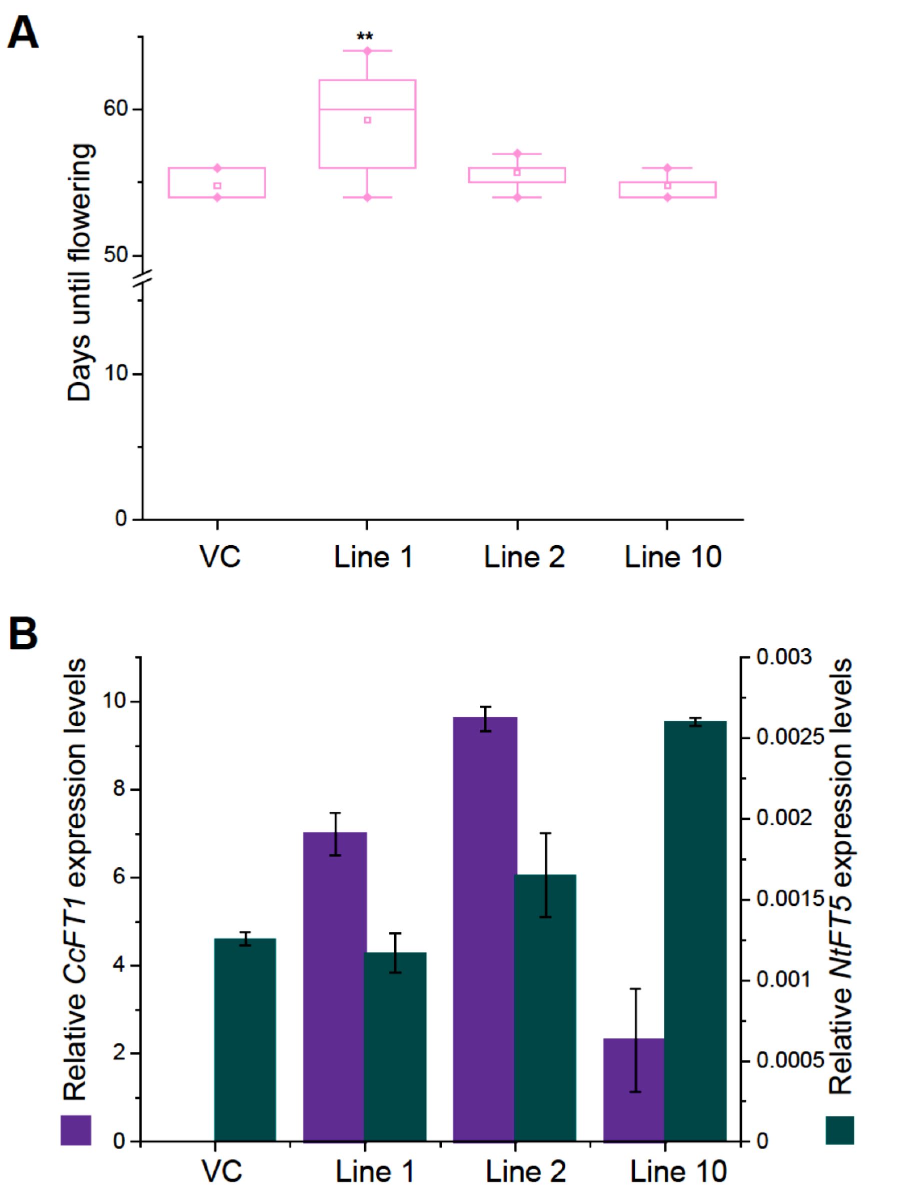


**Fig. 12. Flowering time analysis of tobacco plants overexpressing *CcFT1*. A)** Flowering phenotype of tobacco lines 1, 2, and 10 overexpressing *CcFT1* compared to vector controls (VC). Days until flowering were determined when the first bud had fully opened. **Statistically significant difference as determined by ANOVA and Tukey's *post hoc* test, $p < 0.01$; $n = 10$ (lines 1, 2 and 10), $n = 5$ (VC). In the boxplots, center line = median, square = mean, box limits = upper and lower quartiles, whiskers = 1.5× interquartile range, and diamonds = outliers. **B)** Overexpression levels of *CcFT1* and endogenous expression levels of *NtFT5* in transgenic plants were determined when plants entered the reproductive growth phase. Data are means ± SEM ($n = 3$ biological replicates).

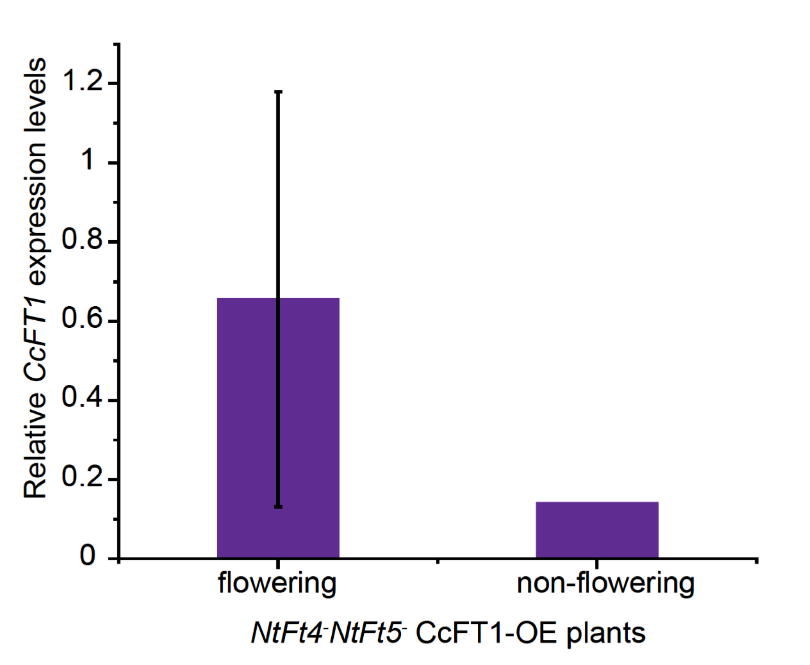


**Fig. 13. Relative expression levels of *CcFT1* in flowering versus non-flowering *Ntft4*⁻*Ntft5*⁻ plants overexpressing *CcFT1*.** Data are means ± SD (*n* = 46 flowering plants and *n* = 2 non-flowering plants).

**Table S1. RNA Seq data of *Cuscuta campestris* and *C. australis*.**

| Organism | Tissue | SRA Accession | Illumina read length, mode | Bases (Gbp) |
|---|---|---|---|---|
| C. australis | Stem | SRR19610423 | 151 bp, PE | 6,445 |
| *C. australis* | Stem | SRR19611357 | 151 bp, PE | 6,505 |
| *C. australis* | Stem | SRR19611520 | 151 bp, PE | 6,386 |
| *C. australis* | Stem | SRR19611521 | 151 bp, PE | 5,863 |
| *C. australis* | Stem | SRR19612874 | 151 bp, PE | 5,699 |
| *C. australis* | Stem | SRR19612875 | 151 bp, PE | 6,616 |
| *C. australis* | Stem | SRR19612878 | 151 bp, PE | 6,220 |
| *C. australis* | Stem | SRR19627926 | 151 bp, PE | 6,231 |
| *C. australis* | Stem | SRR19627939 | 151 bp, PE | 6,261 |
| *C. australis* | Stem | SRR20661662 | 151 bp, PE | 6,186 |
| *C. australis* | Stem | SRR20662442 | 151 bp, PE | 6,547 |
| *C. australis* | Stem | SRR20662443 | 151 bp, PE | 5,744 |
| *C. australis* | Germinating seedling | SRR6664647 | 151 bp, PE | 6,087 |
| *C. australis* | Haustoria on tomato | SRR6664648 | 151 bp, PE | 7,236 |
| *C. australis* | Bud | SRR6664649 | 151 bp, PE | 8,374 |
| *C. australis* | Ovary | SRR6664650 | 151 bp, PE | 8,936 |
| *C. australis* | Stem tip on tomato | SRR6664651 | 151 bp, PE | 7,697 |
| *C. australis* | Seeds | SRR6664652 | 151 bp, PE | 7,329 |
| *C. australis* | Pre-haustoria on tomato | SRR6664653 | 151 bp, PE | 8,503 |
| *C. australis* | Stems on tomato | SRR6664654 | 151 bp, PE | 7,452 |
| | | | | **136,328** |
| *C. campestris* | Not specified* | ERR1916345 | 125 bp, PE | 5,062 |
| *C. campestris* | Not specified* | ERR1916346 | 125 bp, PE | 1,832 |
| *C. campestris* | Not specified* | ERR1916347 | 125 bp, PE | 8,524 |
| *C. campestris* | Not specified* | ERR1916348 | 125 bp, PE | 20,608 |
| *C. campestris* | Not specified* | ERR1916349 | 125 bp, PE | 17,925 |
| *C. campestris* | Not specified* | ERR1916350 | 125 bp, PE | 7,865 |
| *C. campestris* | Not specified* | ERR1916351 | 125 bp, PE | 8,375 |
| *C. campestris* | Not specified* | ERR1916352 | 125 bp, PE | 8,777 |
| *C. campestris* | Not specified* | ERR1916353 | 125 bp, PE | 5,125 |
| *C. campestris* | Not specified* | ERR1916354 | 125 bp, PE | 2,518 |
| *C. campestris* | Not specified* | ERR1916355 | 125 bp, PE | 14,034 |
| *C. campestris* | Not specified* | ERR1916356 | 125 bp, PE | 6,807 |
| *C. campestris* | Not specified* | ERR1916357 | 125 bp, PE | 0.541 |
| *C. campestris* | Not specified* | ERR1916358 | 125 bp, PE | 24,504 |
| *C. campestris* | Not specified* | ERR1916359 | 125 bp, PE | 6,649 |
| *C. campestris* | Not specified* | ERR1916360 | 125 bp, PE | 1,202 |
| *C. campestris* | Not specified* | ERR1916361 | 125 bp, PE | 4,443 |
| *C. campestris* | Not specified* | ERR1916362 | 125 bp, PE | 13,772 |

| | | | | |
|---|---|---|---|---|
| *C. campestris* | Not specified* | ERR1916363 | 125 bp, PE | 5,153 |
| *C. campestris* | Not specified* | ERR1916364 | 125 bp, PE | 3,823 |
| *C. campestris* | Penetrating haustorium | SRR12763776 | 101 bp, PE | 3,121 |
| *C. campestris* | Attaching haustorium | SRR12763777 | 101 bp, PE | 3,094 |
| *C. campestris* | Attaching haustorium | SRR12763778 | 101 bp, PE | 3,175 |
| *C. campestris* | Attaching haustorium | SRR12763779 | 101 bp, PE | 3,168 |
| *C. campestris* | Swelling haustorium | SRR12763780 | 101 bp, PE | 3,181 |
| *C. campestris* | Swelling haustorium | SRR12763781 | 101 bp, PE | 3,194 |
| *C. campestris* | Swelling haustorium | SRR12763782 | 101 bp, PE | 3,187 |
| *C. campestris* | Vegetative stem | SRR12763783 | 101 bp, PE | 3,176 |
| *C. campestris* | Penetrating haustorium | SRR12763787 | 101 bp, PE | 3,130 |
| *C. campestris* | Penetrating haustorium | SRR12763788 | 101 bp, PE | 3,204 |
| *C. campestris* | Vegetative stem | SRR12763789 | 101 bp, PE | 3,119 |
| *C. campestris* | Vegetative stem | SRR12763790 | 101 bp, PE | 3,095 |
| *C. campestris* | Haustorial region | SRR13299802 | 51 bp, SE | 0.387 |
| *C. campestris* | Haustorial region | SRR13299801 | 51 bp, SE | 0.396 |
| *C. campestris* | Haustorial region | SRR13299800 | 51 bp, SE | 0.618 |
| *C. campestris* | Haustorial region | SRR13299799 | 51 bp, SE | 0.714 |
| *C. campestris* | Haustorial region | SRR13299798 | 51 bp, SE | 0.172 |
| *C. campestris* | Haustorial region | SRR13299797 | 51 bp, SE | 0.301 |
| *C. campestris* | Haustorial region | SRR13299796 | 51 bp, SE | 0.212 |
| *C. campestris* | Haustorial region | SRR13299795 | 51 bp, SE | 0.308 |
| *C. campestris* | Haustorial region | SRR13299794 | 51 bp, SE | 0.225 |
| *C. campestris* | Haustorial region | SRR13299793 | 51 bp, SE | 0.283 |
| *C. campestris* | Haustorial region | SRR14853881 | 151 bp, PE | 8,619 |
| *C. campestris* | Haustorial region | SRR14853883 | 151 bp, PE | 8,954 |
| *C. campestris* | Haustorial region | SRR14853885 | 151 bp, PE | 11,372 |
| *C. campestris* | Haustorial region | SRR14853887 | 151 bp, PE | 9,394 |
| | | | | **247,357** |

* Vogel et al. (2018) used "non-infectious feeding stems, detached starving stems and different infection stages", p.8 (2); PE – paired-end mode; SE – single-end mode.

**Table S2. Taxon list and accession numbers for PEBP proteins used in gene family analysis.**

| Species | Gene denomination (if any) | Locus tag, Accession number |
|---|---|---|
| *Arabidopsis thaliana* | ABI5 | AT2G36270 |
| *Arabidopsis thaliana* | AREB3 | AT3G56850 |
| *Arabidopsis thaliana* | FD | AT4G35900 |
| *Arabidopsis thaliana* | FDP (BZIP27) | AT2G17770 |
| *Brachypodium distachyon* | BdFD1 | Bradi3g60870 |
| *Brachypodium distachyon* | BdFD2 | Bradi4g36587 |
| *Brachypodium distachyon* | BdFD3 | Bradi1g29920 |
| *Cuscuta australis* | BZIP-family gene | g21250.t1 |
| *Cuscuta australis* | BZIP-family gene | g56588.t1-CDS |
| *Cuscuta australis* | BZIP-family gene | g2115.t1 |
| *Cuscuta australis* | BZIP-family gene | g29401.t1 |
| *Cuscuta australis* | *CaFD | *C046N0061E0.1 |
| *Cuscuta australis* | CaFD-like | C001N0026E0.1; |
| *Cuscuta australis* | CaFD-like | C025N0100E0.1 |
| *Cuscuta australis* | CaFD-like | C033N0055E0.1 |
| *Cuscuta australis* | CaFD-like | C056N0011E0.1 |
| *Cuscuta australis* | CaFD-like | C036N0249E0.1 |
| *Cuscuta campestris* | CcFD-like | Ccam0.32, scaffold247, Cc032062 |
| *Cuscuta campestris* | CcFD-like | Ccam0.32, scaffold2, Cc010070 |
| *Cuscuta campestris* | CcFD-like | Ccam0.32, scaffold30, Cc002951 |
| *Cuscuta campestris* | FD-like | Ccam0.32, scaffold218, Cc030291 |
| *Cuscuta campestris* | CcFD1 | Ccam0.32, scaffold223, Cc030608 |
| *Cuscuta campestris* | CcFD2 | Ccam0.32, scaffold48, OOIL02004257.1 |
| *Fragaria vesca* | FvFD1 | mrna14556.1-v1.0-hybrid |
| *Fragaria vesca* | FvFD2 | mrna08556.1-v1.0-hybrid |
| *Glycine max* | GmFD1 | Glyma04g02420 |
| *Glycine max* | GmFD2 | Glyma06g02470 |
| *Hordeum vulgare* | HvFD1 | BAJ91167 |
| *Hordeum vulgare* | HvFD2 | BAK04622 |
| *Hordeum vulgare* | HvFD3 | AK249012 |
| *Ipomoea nil* | INIL11 | g05356.t1 |
| *Ipomoea triloba* | ItFD | XP_31117467.1 |
| *Malus domestica* | MdFD1 | MDP0000169473 |
| *Malus domestica* | MdFD2 | MDP0000636541 |
| *Musa acuminata* | MaFD1 | GSMUA_Achr1T02630 |
| *Musa acuminata* | MaFD2 | GSMUA_Achr5T11470_001 |
| *Musa acuminata* | MaFD3 | GSMUA_chr9G24090_001 |
| *Oriza sativa* | n.a. | Os01g0813100 |
| *Oriza sativa* | OsFD1 | Os09g0540800 |

| | | |
|---|---|---|
| *Oriza sativa* | OsFD2 | Os06g0720900 |
| *Oriza sativa* | OsFD3 | Os02g0833600 |
| *Oriza sativa* | OsFD4 | Os08g0549600 |
| *Oriza sativa* | OsFD5 | Os06g0724000 |
| *Oriza sativa* | OsFD6 | Os06g0719500 |
| *Oriza sativa* | TRAB1 | Os08g0472000 |
| *Phoenix dactylifera* | PdFD1 | PDK30s1175071g003 |
| *Piecea glauca* | PgABI5-1 | BT102312.1 |
| *Piecea glauca* | PgABI5-2 | BT110053.1 |
| *Pinus taeda* | PtABI5 | AEK86263.1 |
| *Popolus trichocarpa* | PtFD1 | POPTR_0005s11140.1 |
| *Popolus trichocarpa* | PtFD2 | POPTR_0005s26480 |
| *Setaria italica* | SiFD1 | Si031077m |
| *Setaria italica* | SiFD2 | Si007412m |
| *Setaria italica* | SiFD3 | Si023448m |
| *Setaria italica* | SiFD4 | Si014546m |
| *Solanum lycopersicum* | SlFD2 | Solyc02g061990.2.1 |
| *Solanum lycopersicum* | SPGB | Solyc02g083520 |
| *Solanum tuberosum* | StFD1 | Sotub02g026810 |
| *Solanum tuberosum* | StFD2 | Sotub02g009830 |
| *Sorghum bicolor* | SbFD1 | Sb02g031340 |
| *Sorghum bicolor* | SbFD2 | Sb10g030400 |
| *Sorghum bicolor* | SbFD3 | Sb04g038600 |
| *Triticum aestivum* | TaFD1 | CK206464 |
| *Triticum aestivum* | TaFDL13 | ABZ91908 |
| *Triticum aestivum* | TaFDL15 | ABZ91909 |
| *Triticum aestivum* | TaFDL2 | ABZ91911 |
| *Triticum aestivum* | TaFDL3 | ABZ91912 |
| *Vitis vinifera* | VvFD1 | GSVIVT01009970001 |
| *Vitis vinifera* | VvFD2 | GSVIVT01006332001 |
| *Zea mays* | DLF1 | GRMZM2G067921 |
| *Zea mays* | ZmFD2 | GRMZM2G402862 |
| *Zea mays* | ZmFD3 | GRMZM2G073892 |

*CaFD sequence identified by Shen et al. 2020 (17)

**Table S3. Taxon list and accession numbers of FD-like proteins for phylogenetic inference.**

| Species | Gene denomination (if any) | Locus tag, Accession number |
|---|---|---|
| *Aegilops tauschii* | FT | ABI34864.1 |
| *Antirrhinum majus* | CENTRORADIALIS-like | CAC21563.1 |
| *Antirrhinum majus* | CENTRORADIALIS-like | CAC21564.1 |
| *Aquilegia formosa* | TERMINAL FLOWER 0 | ABB90591.1 |
| *Arabidopsis lyrata* subsp. *lyrata* | CENTRORADIALIS-like | XP_2876387.1 |
| *Arabidopsis lyrata* subsp. lyrata | TERMINAL FLOWER 0 | AAM27957.1 |
| *Arabidopsis lyrata* subsp. *lyrata* | TERMINAL FLOWER 1 | XP_2873117.1 |
| *Arabidopsis lyrata* subsp. *lyrata* | BROTHER of FT and TFL 1 | XP_2864769.1 |
| *Arabidopsis thaliana* | PEBP family protein | NP_176726.1 |
| *Arabidopsis thaliana* | PEBP family protein | NP_193770.1 |
| *Arabidopsis thaliana* | CENTRORADIALIS-like | NP_180324.1 |
| *Arabidopsis thaliana* | PEBP family protein | NP_196004.1 |
| *Arabidopsis thaliana* | PEBP family protein | NP_201010.1 |
| *Arabidopsis thaliana* | PEBP family protein | NP_173250.1 |
| *Beta vulgaris* | flowering locus T-like protein FT2 | ADM92610.1 |
| *Beta vulgaris* | flowering locus T-like protein FT1 | ADM92608.1 |
| *Beta vulgaris* | centroradialis-like protein CEN1 | ADM92611.1 |
| *Brassica napus* | BNTFL1-1 | BAA33415.1 |
| *Brassica napus* | BNTFL1-3 | BAA33417.1 |
| *Brassica napus* | BNTFL1-2 | BAA33416.1 |
| *Brassica rapa* | BRTFL1-1 | BAA33418.1 |
| *Brassica rapa* | BRTFL1-2 | BAA33419.1 |
| *Capsicum annuum* | self-pruning-like protein | AAZ66798.1 |
| *Capsicum frutescens* | fasciculate | ACL27223.1 |
| *Carica papaya* | flowering locus T | ACX85427.1 |
| *Chaenomeles sinensis* | TFL1-like protein | BAD10965.1 |
| *Chaenomeles sinensis* | TFL1-like protein | BAD10971.1 |
| *Chrysanthemum lavandulifolium* | flowering locus T-like protein | ACY82397.2 |
| *Chrysanthemum* x *morifolium* | flowering locus T-like protein | ACX48949.1 |
| *Chrysanthemum* x *morifolium* | flowering locus T-Like protein | BAJ14266.2 |
| *Citrus sinensis* | terminal flower | AAR04683.1 |
| *Citrus sinensis* | terminal flower | AAR04684.1 |
| *Citrus trifoliata* | CTRSFT1-like protein | ABY91244.1 |
| *Citrus trifoliata* | CTRSTFL-like protein | ABY91242.1 |
| *Citrus trifoliata* | CTRSTFL-like protein | ABY91243.1 |
| *Citrus unshiu* | CiFT | BAA77836.1 |
| *Citrus unshiu* | flowering locus T | BAF96644.1 |
| *Citrus unshiu* | flowering locus T | BAF96645.1 |
| *Citrus unshiu* | MOTHER of FT, TFL1-like protein | BAF93494.1 |
| *Crocus sativus* | terminal flower 1-like protein | ACX53295.1 |
| *Cucumis sativus* | FT-like protein | BAH28253.1 |
| *Cucumis sativus* | TFL1-like protein | BAH28255.1 |
| *Cucurbita maxima* | flowering locus T-like 2 | ABI94606.1 |

| | | |
|---|---|---|
| *Cucurbita maxima* | flowering locus T-like 1 | ABI94605.1 |
| *Cucurbita moschata* | FTL2 | ABR20499.1 |
| *Cucurbita moschata* | FTL1 | ABR20498.1 |
| *Cuscuta australis* | FT, partial | DOI: 10.5061/dryad.jsxksn0dr |
| *Cuscuta australis* | C020N0378E0.1 | CaFT |
| *Cuscuta campestris* | CcFT1 | DOI: 10.5061/dryad.jsxksn0dr |
| *Cuscuta campestris* | CcFT2 | DOI: 10.5061/dryad.jsxksn0dr |
| *Cuscuta campestris* | TL1-like partial | DOI: 10.5061/dryad.jsxksn0dr |
| *Cydonia oblonga* | TFL1-like protein | BAD10964.1 |
| *Cydonia oblonga* | TFL1-like protein | BAD10970.1 |
| *Erythrante guttata* | FT, partial | DOI: 10.5061/dryad.jsxksn0dr |
| *Ficus carica* | flowering locus T-like protein | BAI60052.1 |
| *Glycine max* | Dt1 | ADF30893.1 |
| *Glycine max* | brother of FT and TFL1 protein | ABS57463.1 |
| *Glycine max* | CETS1 | ABF65987.1 |
| *Glycine max* | mother of FT-like protein | ACA24491.1 |
| *Glycine max* | unknown | ACU14433.1 |
| *Gossypium arboreum* | terminal flower 1a | ABW24961.1 |
| *Gossypium arboreum* | terminal flower 1b | ABW24962.1 |
| *Gossypium arboreum* | terminal flower 1b | ABW24969.1 |
| *Gossypium hirsutum* | terminal flower 1a | ABW24963.1 |
| *Gossypium hirsutum* | terminal flower 1a | ABW24970.1 |
| *Gossypium hirsutum* | terminal flower 1a | ABW24967.1 |
| *Gossypium hirsutum* | terminal flower 1b | ABY62770.1 |
| *Gossypium hirsutum* | terminal flower 1b | ABW24964.1 |
| *Gossypium raimondii* | terminal flower 1a | ABW24965.1 |
| *Gossypium raimondii* | terminal flower 1b | ABW24966.1 |
| *Helianthus annuus* | flowering locus T4 | ADF32945.1 |
| *Helianthus annuus* | flowering locus T2 | ADF32947.1 |
| *Helianthus annuus* | terminal flower 1 | ADO61015.1 |
| *Hordeum vulgare* | FT | ABI55201.1 |
| *Hordeum vulgare* | FT | ABI55202.1 |
| *Hordeum vulgare* | FT | ABJ97441.1 |
| *Hordeum vulgare* subsp. *vulgare* | vrn-H3 late flowering allele | ABK91684.1 |
| *Hordeum vulgare* subsp. *vulgare* | FT-like protein | ABB99414.1 |
| *Hordeum vulgare* subsp. *vulgare* | terminal flower 1-like protein | ABF85670.1 |
| *Hordeum vulgare* subsp. *vulgare* | homologous protein to TFL1 | BAH24197.1 |
| *Impatiens balsamina* | TERMINAL FLOWER 1 protein | CAI61980.1 |
| *Impatiens balsamina* | TERMINAL FLOWER 1 protein | CAI61982.1 |
| *Ipomoea nil* | FT-like protein | ABW73563.1 |
| *Ipomoea nil* | FT-like protein | ABW73562.1 |
| *Ipomoea nil* | CENTRORADIALIS homolog | BAE44112.1 |
| *Ipomoea triloba* | FT, partial | DOI: 10.5061/dryad.jsxksn0dr |
| *Lactuca sativa* | FT, partial | DOI: 10.5061/dryad.jsxksn0dr |
| *Leavenworthia crassa* | terminal flower 1-like protein | ADC32542.1 |
| *Lolium perenne* | FT3 | ABC33722.1 |

| | | |
|---|---|---|
| *Lolium perenne* | terminal flower 1-like protein | AAG31808.1 |
| *Lotus japonicus* | CEN/TFL1-like protein | AAQ93599.1 |
| *Malus domestica* | flowering locus T | ACL98164.1 |
| *Malus domestica* | flowering locus T like protein | BAD08340.1 |
| *Malus domestica* | CENTRORADIALIS like protein | BAG31957.1 |
| *Malus domestica* | CENTRORADIALIS like protein | BAG31958.1 |
| *Malus domestica* | TFL1-like protein | BAD06418.1 |
| *Malus domestica* | TFL1-like protein | BAD10961.1 |
| *Malus domestica* | TFL1-like protein | BAD10967.1 |
| *Malus domestica* | TFL1 like protein | BAG31959.1 |
| *Medicago truncatula* | PEBP | ABE80135.1 |
| *Misopates orontium* | CEN/TFL1-like protein | CAJ44126.1 |
| *Nicotiana tabacum* | NtFT5 | DOI: 10.5061/dryad.jsxksn0dr |
| *Nicotiana tabacum* | NtFT4 | DOI: 10.5061/dryad.jsxksn0dr |
| *Nicotiana tabacum* | CEN-like protein 2 | AAD43529.1 |
| *Nicotiana tabacum* | CEN-like protein 4 | AAD43530.1 |
| *Nicotiana tabacum* | CEN-like protein 1 | AAD43528.1 |
| *Olea europaea* | FT, partial | DOI: 10.5061/dryad.jsxksn0dr |
| *Oncidium* hybrid cultivar | flowering locus T | ACC59806.1 |
| *Oryza glumipatula* | FT-like protein | BAH56285.1 |
| *Oryza longistaminata* | FT-like protein | BAH30248.1 |
| *Oryza rufipogon* | Hd3a | BAG72302.1 |
| *Oryza rufipogon* | FT-like protein | BAH30250.1 |
| *Oxybasis rubra* | Flowering locus T-like 1 protein | ABV56568.1 |
| *Oxybasis rubra* | flowering locus T-like 2 | ABP02016.1 |
| *Phyllostachys meyeri* | FT-like protein | BAI49899.1 |
| *Phyllostachys meyeri* | FT-like protein | BAI49900.1 |
| *Phyllostachys meyeri* | FT-like protein | BAI49901.1 |
| *Picea abies* | FT-like/TFL1-like protein | ABQ85553.1 |
| *Picea sitchensis* | unknown | ABK25607.1 |
| *Picea sitchensis* | unknown | ACN40175.1 |
| *Pisum sativum* | TFL1a | AAR03725.1 |
| *Populus deltoides* | flowering locus T-like protein FT1 | AAS00056.1 |
| *Populus nigra* | flowering locus T | BAD01561.1 |
| *Populus nigra* | flowering locus T | BAD01612.1 |
| *Populus nigra* | flowering locus T | BAD02371.1 |
| *Populus nigra* | FLOWERING LOCUS T | BAG12904.1 |
| *Populus nigra* | terminal flower 1 | BAD22599.1 |
| *Populus nigra* | flowering locus T like protein | BAD22601.1 |
| *Populus nigra* | flowering locus T like protein | BAD27481.1 |
| *Populus nigra* | flowering locus T like protein | BAD22677.1 |
| *Populus tremula* | flowering locus T-like protein FT1 | ABD52003.1 |
| *Populus trichocarpa* | protein HEADING DATE 3A | XP_2316173.1 |
| *Populus trichocarpa* | protein HEADING DATE 3A | XP_2311264.1 |
| *Populus trichocarpa* | CEN-like protein 2 | XP_2312811.1 |
| *Populus trichocarpa* | CEN-like protein 1 | XP_2321903.2 |

| | | |
|---|---|---|
| *Populus trichocarpa* | MFT-like protein | ABC26020.1 |
| *Prunus mume* | flowering locus T like protein | CAQ16124.1 |
| *Prunus mume* | TFL1-like protein | BAJ14521.1 |
| *Prunus persica* | flower locus T | ACH73165.1 |
| *Pyrus communis* | TFL1-like protein | BAD10963.1 |
| *Pyrus communis* | TFL1-like protein | BAD10969.1 |
| *Pyrus pyrifolia* | TFL1-like protein | BAD10962.1 |
| *Pyrus pyrifolia* | TFL1-like protein | BAD10968.1 |
| *Raphanus sativus* | terminal flower1-like protein | BAH37010.1 |
| *Rhaphiolepis bibas* | TFL1-like protein | BAD10966.1 |
| *Rhaphiolepis bibas* | TFL1-like protein | BAD10972.1 |
| *Sinapis alba* | flowering locus T | ACM69283.1 |
| *Solanum lycopersicum* | self-pruning protein | AAC26161.1 |
| *Solanum tuberosum* | terminal flower 1 protein | ABC24691.1 |
| *Sorghum bicolor* | protein HEADING DATE 3A | XP_2436509.1 |
| *Sorghum bicolor* | protein TWIN SISTER of FT | XP_2443085.1 |
| *Sorghum bicolor* | protein FLOWERING LOCUS T | XP_2454134.1 |
| *Sorghum bicolor* | CEN-like protein 2 | XP_2442808.1 |
| *Sorghum bicolor* | CEN-like protein 2 | XP_2450283.1 |
| *Sorghum bicolor* | SELF-PRUNING | XP_2453931.1 |
| *Sorghum bicolor* | FLOWERING LOCUS T | XP_2438551.1 |
| *Sorghum bicolor* | FLOWERING LOCUS T | XP_2446272.2 |
| *Sorghum bicolor* | FLOWERING LOCUS T | XP_2451827.2 |
| *Sorghum bicolor* | HEADING DATE 3A | XP_2456354.1 |
| *Sorghum bicolor* | MOTHER of FT/TFL1 homolog 2 | XP_2457494.1 |
| *Spinacia oleraceae* | FT, partial | DOI: 10.5061/dryad.jsxksn0dr |
| *Triticum aestivum* | putative kinase inhibitor | BAG12867.1 |
| *Triticum aestivum* | flowering locus T | AAW23034.1 |
| *Triticum aestivum* | flowering locus T | ACA25439.1 |
| *Triticum aestivum* | flowering locus T, partial | ACA25438.1 |
| *Triticum monococcum* | VRN3 | ABK32206.1 |
| *Triticum turgidum* | VRN3 | ABK32207.1 |
| *Vaccinum darrowii* | FT, partial | DOI: 10.5061/dryad.jsxksn0dr |
| *Vicia faba* | TFL1, partial | ABP73380.1 |
| *Vigna unguiculata* | terminal flower 1a | BAJ22384.1 |
| *Vigna unguiculata* | terminal flower 1b | BAJ22383.1 |
| *Vitis mustangensis* | terminal flower 1 | ADF80900.1 |
| *Vitis vinifera* | flowering locus T-like protein | ABF56526.1 |
| *Vitis vinifera* | FT-like protein | ABI99465.1 |
| *Vitis vinifera* | flowering locus T | ABL98120.1 |
| *Vitis vinifera* | TFL1A protein | ABI99466.1 |
| *Vitis vinifera* | TFL1B protein | ABI99467.1 |
| *Vitis vinifera* | TFL1C protein | ABI99468.1 |
| *Vitis vinifera* | MFT-like protein | ABI99469.1 |

**Table S4. List of primers used in this study.** Restriction sites in 5´-overhangs are underlined. Annealing temperature (T) are given only for PCR primers.

| Primer name | Sequence (5´to 3´) | T [°C] | purpose |
|---|---|---|---|
| Q35S-P SmaI fw | AGACCCGGGTCCAATCCCACCAAAAC | 58 | Generation of pLab12.10Q35S-P |
| Q35S-P SOE rev | GCTCTTATACTTGAGCGTGTCCT | | |
| TMV Ω TL SOE fw | AGGACACGCTCAAGTATAAGAGC | 58 | |
| TMV Ω TL XhoI rev | AGACTCGAGATGGATAATTGTAAATGTAATTGT AATG | | |
| CcFT1-like NcoI fw | AGACCATGGCGGCGAGGCGGCAGA | 68 | Cloning of pLab12.10Q35S-P::*CcFT1* |
| CcFT1-like XbaI rev | AGATCTAGACTATTGACGGCGCCTGCCGC | | |
| CcFT2 PspXI fw | AGACTCGAGCA ACAATGGCGGCGAGGCGGCAGA | 68 | Cloning of pLab12.1Q35S-P::*CcFT2* |
| CcFT2-like XbaI rev | AGATCTAGACTATTAACGGCGCCTGCCTCCG | | |
| CAS9 fw | ACGTGACCGAGGGAATGAGG | 62 | Analysis of CRISPR transgene integration into the genome of *Ntft5*$^{-}$ plants |
| CAS9 rev | TTGCAGGAGATCCAGCGAGG | | |
| NtFT4 ExI CRISPR T3 f | ATTGAAGGTCTGTTGATCTTAGAG | - | *NtFT4* protospacer sequence |
| NtFT4 ExI CRISPR T3 r | AAACCTCTAAGATCAACAGACCTT | - | |
| NtFT4 5'UTRf | GAAAAGTAAAGTATTTAGTGTATAG | 48 | Amplification of *NtFT4* (exon I) for sequencing (4) |
| NtFT4 Intr1 rev | ATGCGAATCAATTATAAATGG | | |
| NtFT5 5'UTR for2 | CCCTTAGACTTGTAAAAACATGC | 53 | Amplification of *NtFT5* (exon I) for sequencing (4) |
| NtFT5 Intr2 rev3 | ATTGTAGATAGAGTTCCTAGCG | | |
| CcFT-like NcoI fw | AGACCATGGCGGCGAGGCGGCAGA | 68 | Cloning of BIFC constructs |
| CcFT1-like XbaI rev | AGATCTAGACTATTGACGGCGCCTGCCGC | | |
| CcFT-like NcoI fw | AGACCATGGCGGCGAGGCGGCAGA | 68 | |
| CcFT2-like XbaI rev | AGATCTAGACTATTAACGGCGCCTGCCTCCG | | |
| CcFD-like NcoI fw | AGACCATGGGATCTCAGGGCGGTGGT | 61 | |
| CcFD-like NheI rev | AGAGCTAGCTCAAAAAGGATTCGAGCTTGTTC | | |
| CcFT ATG fw | ATGGCGGCGAGGCGGCAGA | 68 | Identification of *CcFT1* genomic sequence |
| CcFT ExonII rev | CACTTGGACTTGGAGCATCAGGATC | | |
| CcFT ExonII fw | GTGATGGTGGATCCTGATGCTC | 63 | |
| CcFT1 gDNA1 II rev | TTTAAGTGGGCATTGATTGTTGGTGTCA | | |
| CcFT1 gDNA2 fw | TGACACCAACAATCAATGCC | 61 | |
| CcFT1 gDNA2 rev | TCGTCTGGAAGTCCATACCCAG | | |
| CcFT1 gDNA3 fw | TCCAGACGAACAATCGTGTTA | 59 | |
| CcFT1 gDNA3 rev | AAATTAGGAGTGACGTATTCTTAT | | |
| CcFT1 gDNA4 fw | TGGATAATCGATCTTTTAGAGTCA | 59 | |
| CcFT1 gDNA4 rev | TCTGTGATACTGTGATGCA | | |
| CcFT1 gDNA5 fw | AAAAGTTAGGAGAACACCAAAT | 60 | |
| CcFT1 gDNA5 rev | TATACCTTTCCACTTGATGACA | | |

| | | | |
|---|---|---|---|
| CcFT1 gDNA6 fw | AGTAGCACTTTTAGTTGAGTCA | 59 | |
| CcFT1 gDNA6 rev | AAGTCAAACTGTGACTATCTT | | |
| CcFT1 g_gap fw | GTGTCCAAGTCATTCTCGGATC | 62 | |
| CcFT1 g_gap rev | CACCTTTGCCGCAAGTAGTTG | | |
| CcFT1 gDNA8 fw | GAAGTTCTCTGAACTCACACAAG | 59 | |
| CcFT1 gDNA8 rev | GATAGAAATTAGTTGATAACATTAGGT | | |
| CcFT1 gDNA9 fw | ACCTAATGTTATCAACTAATTTCTATC | 59 | |
| CcFT1 gDNA9 rev | TTTAGTTTTGAACTCTTAAACCTTAAA | | |
| CcFT1 gDNA10 fw | TTTAAGGTTTAAGAGTTCAAAACTAAA | 61 | |
| CcFT1 gDNA10 rev | AGACCCGGAGCAAATAGAT | | |
| CcFT1 gDNA11 fw | GGTCTGGACCTATGGTCTG | 61 | |
| CcFT1 gDNA11 rev | AGTTCACATTGACTCTATGAGTT | | |
| CcFT1 gDNA12 fw | AACTCATAGAGTCAATGTGAACT | 59 | |
| CcFT1 gDNA12 rev | CCTCACACTAAGACATTATTAAG | | |
| CcFT ATG fw | ATGGCGGCGAGGCGGCAGA | 68 | Identification of *CcFT2* genomic sequence |
| CcFT ExonII rev | CACTTGGACTTGGAGCATCAGGATC | | |
| CcFT ExonII fw | GTGATGGTGGATCCTGATGCTC | 64 | |
| CcFT2-IntII_rev | GGAGAGGGATTTAAGGAGAGTGTA | | |
| CcFT2 gDNA1 fw | TACCCATCACATCAAAGGAC | 59 | |
| CcFT2 gDNA1 rev | CATAACTCATTTCACACTTGTC | | |
| CcFT2 gDNA2 fw | GGCATAATGAACGAATTCGAC | 59 | |
| CcFT2 gDNA2 rev | GTCTTAGTGGAAGGAATAATGC | | |
| CcFT2 gDNA3 fw | CCCGTTCCGAATTAATCCTA | 58 | |
| CcFT2 gDNA3 rev | ACCTTTGCCGCAAGTAGTTC | | |
| CcFT2 gDNA4 fw | AGTCAAGCGTGCTTGACTTGAAG | 64 | |
| CcFT2 gDNA4 rev | TGCTATGAGAATCCTAGAACTAGCCA | | |
| CcFT2 gDNA5 fw | CACGTGAGAAGGGACATG | 59 | |
| CcFT2 gDNA5 rev | TCTGCTCGTGTACGATTACAG | | |
| CcFT2 gDNA6 fw | CATACAGAGTATATCTCAATCCTAGA | 59 | |
| CcFT2 gDNA6 rev | CCAAAATTAACTCCGGTGCTCG | | |
| CcFT2 g_gap1 fw | CTTTCTGACCAAGTACGCCAC | 62 | |
| CcFT2 g_gap1 rev | CGTTCAGCCACAGACTACCTACAG | | |
| CcFT2 g_gap2 fw | CACCCAGGAAGACACAGTAACC | 62 | |
| CcFT2 g_gap2 rev | GTCAGTTGGGAAAGCATGTGC | | |
| CcFT2 g_gap3 fw | GGATTGCCATAAGTCTGTATGC | 62 | |
| CcFT2 g_gap3 rev | GCTGGAACAGTACAAATACGAACC | | |
| CcFT2 gap2 seq fw | CTGCATCAGAATAAGCCTGTAG | - | |
| CcFT2 gap2 seq rev | GTAGTTTTCGACGAGCACAC | - | |
| M13 fw | GTAAAACGACGGCCAG | - | Sequencing of subcloned PCR products |
| M13 bw | CAGGAAACAGCTATGAC | - | |
| 3′-RACE | GACTCGTGTGGACATCGATTTTTTTTTTTTTTTTT | - | 3′-RACE of *CcFT* |
| 3′-Adap_1 | GACTCGTGTGGACATCG | 64 | |
| CcFT GSP1 fw | TGCTATGAAAGCCCGCAACC | | |
| 3′-Oligo_d(T) | GCTGTCAACGATACGCTACGTAACGGCATGACA GTGTTTTTTTTTTTTTTTTTTTTTTTT | - | 3′-RACE of *CaFT* |

| | | | |
|---|---|---|---|
| 3′-Adap_2 | GCTGTCAACGATACGCTACGTAACG | 64 | |
| CcFT GSP1 fw | TGCTATGAAAGCCCGCAACC | | |
| 3′-Adap_nested | CGCTACGTAACGGCATGACAGTG | 60 | |
| CcFT GSP2 fw | TCGTACTGTTCCAGCAGATG | | |
| CcFT ATG fw | ATGGCGGCGAGGCGGCAGA | 64 | Amplification of *CcFT1*/*CcFT2*/*CaFT* cDNA |
| 3′-UTR CcFT rev | GCGCACGGTGGATTATTGTTCT | | |
| qPCR CcActin fw | ATGGAAGCTGCTGGAATCCAC | 62 | Expression analysis of *Cuscuta* spp. *Actin* by qRT-PCR (10) |
| qPCR CcActin rev | TTGCTCATACGGTCAGCGATG | | |
| qPCR CaEF1α fw | TCAGACTGTTGCTGTGGGTG | 62 | Expression analysis of *Cuscuta* spp. *EF1a* by qRT-PCR (11) |
| qPCR CaEF1α rev | CCCTTGTGCGGTTCACTTCT | | |
| qPCR CcFT1 disc fw | GGAGAGGTGCTTGAGCCT | 62 | Expression analysis of *CcFT1* and *CaFT* by qRT-PCR |
| qPCR CcFT1 disc rev | CGGGCTTTCATAGCATACC | | |
| qPCR CcFT2 disc fw | GTGATCGGAGAGGTGCTG | 62 | Expression analysis of *CcFT2* by qRT-PCR |
| qPCR CcFT2 disc rev | CTGTGGGCTTTCATAACATACA | | |
| qPCR CcFD fw | ACCGATGACCATGTCTCCAT | 62 | Expression analysis of *CcFD*-like and *CaFD* by qRT-PCR |
| qPCR CcFD rev | CTTGCTCTCAAGCTCCTGTG | | |
| qRTNt/Ntom EF-1α for | AAGCTGACTGTGCTGTCCTGA | 67 | Expression analysis of *NtEF1α* by qRT-PCR (5) |
| qRTNt/Ntom EF-1α rev | GGTGGTAGCATCCATCTTGTTG | | |
| qRTNtFT5/NtomFTy for discr | TCCCAAGTTATTAACCAGCCC | 67 | Expression analysis of *NtFT5* by qRT-PCR (5) |
| qRTNtFT5/NtomFTy rev discr | CATAGCACACAATTTCCTGGC | | |
| qPCR GmF-Box fw | AGATAGGGAAATGGTGCAGGT | 62 | Expression analysis of *GmF-Box* by qRT-PCR (12) |
| qPCR GmF-Box rev | CTAATGGCAATTGCAGCTCTC | | |

**Dataset S1 (separate file). Results of exonerate-based genic conservation analysis of 295 flowering time regulating genes in *Cuscuta* spp. and 13 related eudicots.** Gene and reading frame conservation is provided as length identity score from exonerate-based *protein2genome* alignments, using validated reference proteins of *Arabidopsis thaliana* as queries. In addition to genomic data searches of *Cuscuta* spp., we also examined transcriptome assemblies, the results of which given in separate columns. Finally, we computed the divergence of *Cuscuta's* flowering gene conservation relative to that of the mean length identity score of the here included 13 non-parasitic eudicots, whereby values >1 indicate stronger conservation level and/or longer reading frames in *Cuscuta* spp. than the average, and values <1 suggest less conservation and/or shorter genes in the parasites. Datasets with recovered genic fragments are published in reusable fasta files in the Dryad Data Repository, https://doi.org/10.5061/dryad.jsxksn0dr.

**Dataset S2 (separate file). Domain-based annotation of transposable elements (DANTE) and inferred TE fragments.** Filtered and unfiltered database hits of DANTE searches within the extended CaFT, CcFT1, and CcFT2 gene regions are summarized by length, identity, similarity, and number of interruptions. The retrieved and reference TE domains are provided for every hit, indicating interruptions such as gaps and stop codons by backslashes and asterisks, respectively. Datasets with recovered genic fragments are published in reusable fasta files in the Dryad Data Repository, https://doi.org/10.5061/dryad.jsxksn0dr.
